# A plant-specific clade of serine/arginine-rich proteins regulates RNA splicing homeostasis and thermotolerance in tomato

**DOI:** 10.1101/2024.05.15.594399

**Authors:** Remus RE Rosenkranz, Stavros Vraggalas, Mario Keller, Srimeenakshi Sankaranarayanan, François McNicoll, Karin Löchli, Daniela Bublak, Moussa Benhamed, Martin Crespi, Thomas Berberich, Christos Bazakos, Michael Feldbrügge, Enrico Schleiff, Michaela Müller-McNicoll, Kathi Zarnack, Sotirios Fragkostefanakis

## Abstract

High temperatures cause heat stress (HS), which has negative effects on plant growth and development and affects many cellular processes including pre-mRNA splicing. In tomato plants the splicing profile of many of genes is altered under HS, including that of *HSFA2*, a central transcriptional regulator of thermotolerance. To identify the core splicing regulators of HS-sensitive alternative splicing, we used *HSFA2* as bait and identified two plant-specific members of the serine/arginine-rich family of splicing factors, namely RS2Z35 and RS2Z36, that inhibit *HSFA2* intron splicing. Single and double CRISPR mutants of these proteins show dysregulated splicing of many genes and exhibit lower basal and acquired thermotolerance. Individual-nucleotide resolution UV cross-linking and immunoprecipitation (iCLIP) of tomato leaves revealed that the majority of HS-sensitive alternatively spliced RNAs are bound by RS2Z35 and RS2Z36 and this interaction occurs at purine-rich RNA motifs. Phenotypic and transcriptome analyses revealed that RS2Z35 and RS2Z36 are important players in the stress response and thermotolerance in plants that mitigate the negative effects of HS on RNA splicing homeostasis.

## Introduction

High temperatures can cause heat stress (HS) and thereby have negative effects on plant growth and development. Survival and recovery from HS are dependent on the synthesis of proteins, like heat shock proteins (HSPs) and reactive oxygen species (ROS) scavengers that protect cellular structures and other proteins from irreversible damages ^1^. The induction of many HSPs but also other HS-responsive genes is controlled by HS transcription factors (HSFs) ^2,3^. In plants, members of the HSFA1 subfamily are essential for the initiation of the HS response and have, therefore, been characterized as master regulators of thermotolerance ^4,5^. The activity of HSFA1 proteins is reinforced by interaction with other HSFs, such as HSFA2 and HSFA7 ^6–8^. These hetero-oligomeric complexes provide the strong transactivation activity required for the upregulation of many genes which are important for acclimation and survival of plants under severe HS conditions ^7,9–11^.

While many studies have focused on transcription as a pivotal step for the regulation of gene expression under HS, pre-mRNA splicing, a crucial step for the synthesis of mRNAs from intron-containing eukaryotic genes, has attracted less attention. Splicing is a temperature-sensitive process and consequently hundreds of genes are alternatively spliced under HS ^12,13^. Alternative splicing affects both constitutively and stress-responsive genes and results in the generation of either aberrant mRNAs or mRNA variants that are translated into alternative protein isoforms. A global inhibition of pre-mRNA splicing is associated with mRNA degradation by the nonsense-mediated mRNA decay (NMD) mechanism, although some intron-containing mRNAs are resistant to NMD, and can be post-transcriptionally spliced before their release to the cytoplasm ^14^. It is assumed, that these mechanisms contribute to stress adaptation and cellular survival by protecting the cell from the accumulation of deleterious protein species and by reducing protein synthesis under proteotoxic conditions ^15^. In addition, alternative splicing contributes to the generation of splice variants that code for protein isoforms with distinct properties and functions. Characteristic examples are *HSFA2* and *HSFA7* in tomato ^7,16^. Splicing of the second intron of these genes results in protein isoforms that are rapidly degraded, while full or partial intron retention yields mRNAs that are translated to stable protein isoforms that are maintained for a long period in thermotolerant cells. Therefore, in the case of tomato but also other Solanaceae species, alternative splicing of the *HSFA2* and *HSFA7* mRNAs is important for acquired thermotolerance (ATT) ^7,16^.

In contrast to the well-established knowledge on the core transcription factors regulating the HS response, the main regulators of HS-sensitive alternative splicing are yet to be found. Some studies have shown that specific splicing factors such as splicing factor 1/branchpoint binding protein (SF1/BBP), LSM5 and STABILIZED1 (STA1) are involved in the splicing of Arabidopsis HSFs and therefore contribute to thermotolerance ^17–19^. Serine/Arginine-rich splicing factors (SRSFs) are involved in both constitutive and alternative splicing ^20^ and members of this family are involved in abiotic stress tolerance ^21^, but their relevance in HS-mediated alternative splicing has not been explored so far.

Here, we find that a 1-hour exposure of tomato leaves to 40°C triggers a strong HS response and alters the splicing profile of a high number of genes that are either differentially or constitutively expressed. We sought to identify the core regulators of HS-sensitive alternative splicing by using a prey-bait approach which included SR proteins as splicing effectors (prey) and *HSFA2* as a potential target (bait). Two proteins that belong to the plant-specific RS2Z subfamily bind to *HSFA2* mRNA and promote intron retention. Phenotypic and transcriptome analysis of single and double *rs2z* CRIPSR mutants and individual-nucleotide resolution UV cross-linking and immunoprecipitation (iCLIP) of tomato leaves showed that RS2Z35 and RS2Z36 are core regulators of RNA splicing and thermotolerance, both in a *HSFA2*-dependent and independent manner.

## Results

### Heat stress affects alternative pre-mRNA splicing in tomato leaves

To elucidate the effect of HS on alternative splicing in tomato, an RNA-seq analysis on leaves exposed to 40°C or 25°C for 1 hour was performed. This treatment triggers a strong HS response evidenced by the induction of key thermotolerance marker genes such as HSPs ^7^. HS resulted in a total of 4785 differentially regulated alternative splicing events (DAS; absolute delta percent selected index [|ΔPSI|] ≥ 0.05, probability changing [*P*_(|ΔPSI| ≥ 0.02)_] ≥ 0.9; Supplementary Dataset 1). The majority of these are binary splicing events (bDAS), namely intron retention (IR), alternative 3’splice selection (A3ss), cassette exon (CE) and alternative 5’ splice selection (A5ss) (Fig. 1a). Altogether, we identified 2801 genes with bDAS events (Supplementary Dataset 1). HS causes both inhibition and stimulation of alternative splicing (Fig. 1b), indicating that the selected treatment is not inhibitory for splicing but rather disturbs the homeostatic balance of RNA splicing. Based on this, we assumed the presence of splicing factors that actively mediate HS-sensitive alternative splicing in a positive and/or negative manner.

**Figure 1.**
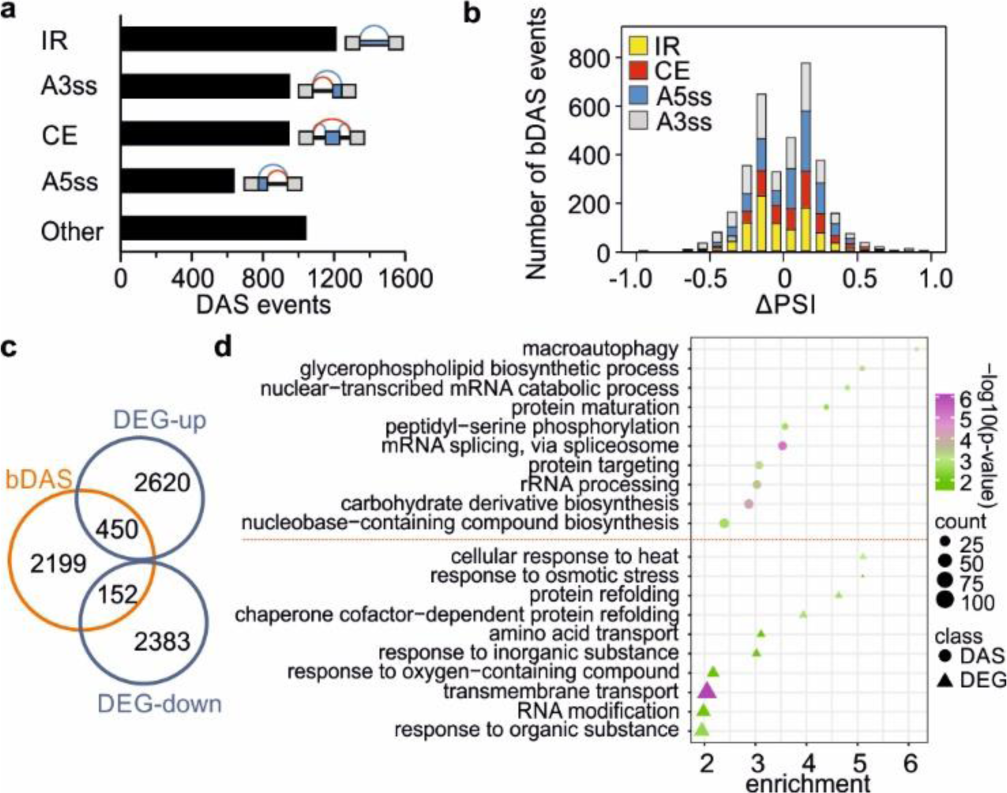
Transcriptome regulation by differential expression and RNA splicing in response to heat stress (40°C, 1 h). (a) Number of differential alternative splicing events (DAS; *P*_(|ΔPSI| ≥ 0.02)_ ≥ 0.9, |ΔPSI| ≥ 0.05). IR: intron retention; A3ss: alternative 3′ splice site; CE: cassette exon; A5ss: alternative 5′ splice site. For details see Supplementary Dataset 1. (b) Distribution of DAS binary (bDAS) events of the IR, CE, A3ss and A5ss type. ΔPSI values indicate enhanced or reduced splicing. (c) Overlap of DAS and differentially expressed genes (DEGs; |log_2_FC| > 1, *p_adj_* < 0.01). (d) Gene ontology (GO) annotation of biological processes of genes identified as DAS or DEG. The *p*-value is calculated based on Fischer’s exact test and Bonferroni correction for multiple testing using the PANTHER GO database. Only the top 10 categories are shown. The complete list is provided in Supplementary Dataset 3.

Among the 2801 bDAS genes, 450 are up- and 152 are downregulated in response to HS, which represent only a small fraction of the total 5605 differentially expressed genes (DEGs; absolute log_2_-transformed fold change [|log_2_FC|] > 1, adjusted *p*-value [*P_adj_*] < 0.01); Fig. 1c; Supplementary Dataset 2). Within the 450 bDAS/DEG genes that are transcriptionally upregulated, we identified *HSFA2* a central factor of ATT, but also genes involved in protein folding and fate, as well as in pre-mRNA splicing regulation ^9,16^. Gene ontology (GO) enrichment analysis revealed that bDAS genes are involved in 33 biological processes while DEGs in only 11, with no overlap between them (Fig. 1d; Supplementary Dataset 3). DEGs are mainly involved in stress responses, while bDAS genes cover a broader spectrum of biological processes including mRNA splicing.

### RS2Z-mediated regulation of *HSFA2* splicing

*HSFA2* is a HS-induced and alternatively spliced gene (Fig. 2a). Full or partial retention of intron 2 results in three splice variants, which all code for the protein isoform HsfA2-I, involved in ATT ^16^. Intron splicing yields isoform HsfA2-II which is rapidly degraded and therefore plays no significant role in thermotolerance of cultivated tomato accessions ^16^. Based on its importance for the HS response, *HSFA2* was used as a bait for the identification of putative regulators of HS-sensitive alternative splicing events (Fig. 2b). Members of the family of SRSFs, which are core regulators of alternative splicing in eukaryotes, were used as prey ^22^. SRSFs in plants are divided into six subfamilies, including three that belong to plant-specific clades ^23^. Seventeen canonical SRSF genes have been identified so far in the tomato genome ^24^. The splicing profile of the endogenous *HSFA2* was examined in heat-stressed tomato mesophyll protoplasts (37.5°C / 1 h) transfected with a plasmid carrying a PCaMV35S::HA-SRSF gene expression cassette (Fig. 2b; Supplementary Fig. S1). The splicing profile of *HSFA2* in protoplasts expressing each SRSF was examined by RT-PCR and compared to a mock control (no transgene SRSF expressed). Five SRSFs, namely RS30, SCL29, SC30b, RS2Z35 and RS2Z36, reduced intron splicing in *HSFA2* while the others had no effect (Fig. 2c).

**Figure 2.**
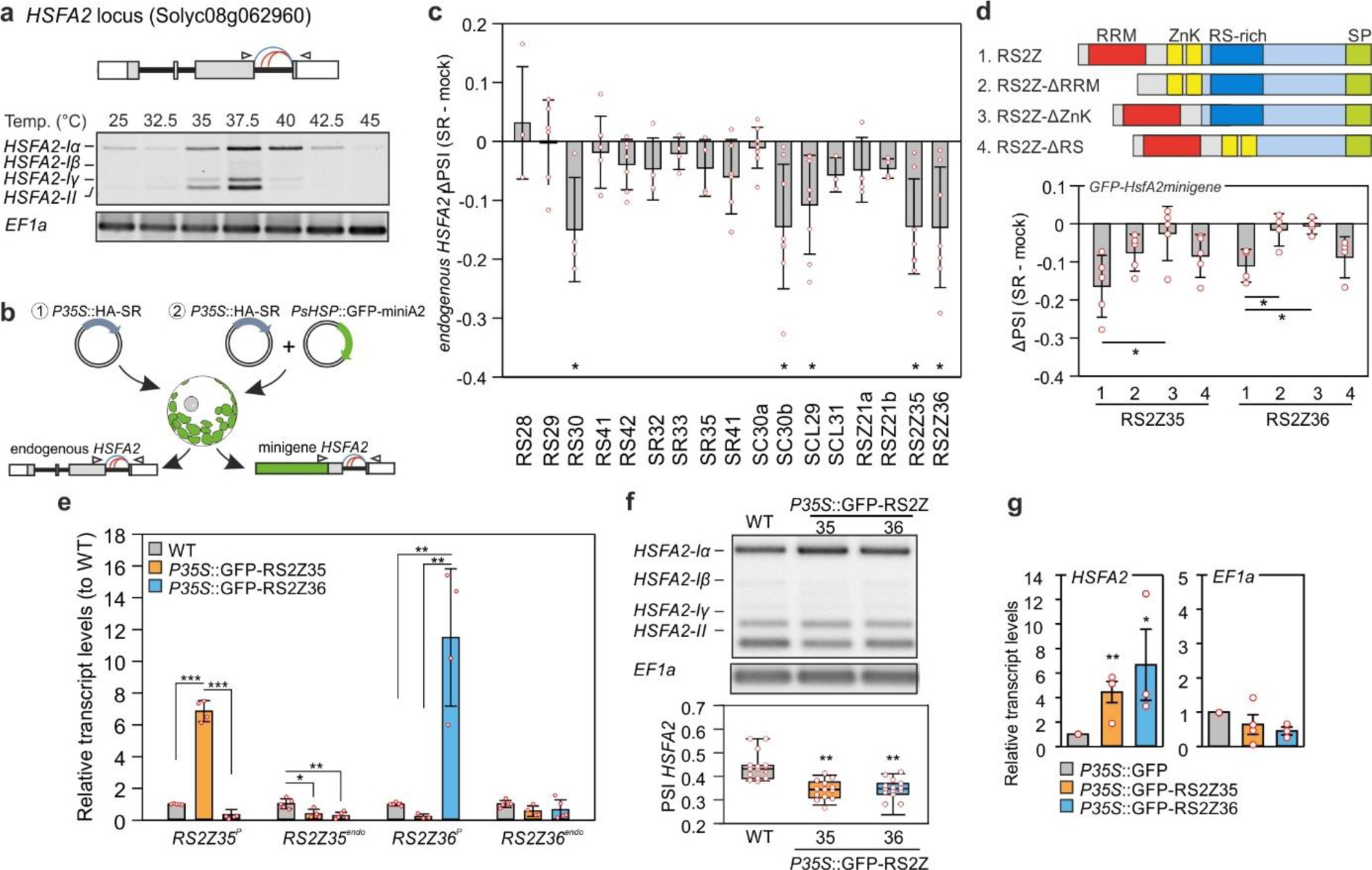
Regulation of *HSFA2* splicing by RS2Z proteins. (a) Splicing profile of *HSFA2*. Top: *HSFA2* exon/intron gene structure. Blue and red curves show canonical and alternative splice sites, respectively. Bottom: RNA-Seq read coverage for *HSFA2*. Splicing profile of *HSFA2* based on semi-quantitative PCR analysis in protoplasts exposed to the indicated temperatures for 1 hour. Oligonucleotides used for PCR anneal in exon 2 and 3 and are depicted by arrowhead in the gene scheme. (b) Strategies used to identify regulators of *HSFA2* splicing based on transient expression of HA-tagged proteins in tomato mesophyll protoplasts. 1: splicing analysis on the endogenous; 2: splicing analysis based on *PHSP21.5::GFP-HSFA2minigene*. (c) Difference in splicing index (PSI) of the endogenous *HSFA2* between SR-expressing and mock protoplasts. Asterisks indicate statistically significant difference (*p* < 0.05) between the SR-expressing and mock sample. Bars are the average of at least three independent biological replicates, ± standard error of mean (SEM). (d) Effect of RS2Z35 and RS2Z36 proteins and domain deletion mutants on splicing of the *HSFA2* minigene driven by the HS-inducible HSP21.5 promoter. Protoplasts were exposed to 37.5°C for 1 hour. Bars are the averages of 3-4 independent experiments. (e) Relative transcript levels of endogenous (endo) and protein-coding (P) *RS2Z35* and *RS2Z36* in WT, and *PCaMV35S::GFP-RS2Z* transgenic lines, based on qRT-PCR analysis. Asterisks indicate statistically significant difference (\**p* < 0.05, \*\**p* < 0.01, \*\*\**p* < 0.001) between the indicated samples. Bars are the average of 3 independent biological replicates, ±SEM. (f) Splicing profile of *HSFA2* in WT and GFP-RS2Z transgenic lines. The median is indicated in the box, with the lower upper quartiles, margined by the extreme values within the interval of 1.5 the interquartile from the box (whiskers). Asterisks shows significant statistical difference (\*\**p* < 0.01) between WT and transgenic lines based on ANOVA and Duncan’s Multiple Range test based on 14 independent experiments. (g) Validation of association of RS2Z proteins with endogenous *HSFA2* or *EF1a* mRNAs via RNA immunoprecipitation (RIP)-qPCR. Bars are the average of 3-4 biological replicates, ±SEM. Asterisks indicate statistically significant difference (\**p* < 0.05, \*\**p* < 0.01, pairwise *t*-test) between RIP samples from the GFP-RS2Z and GFP transgenic lines.

We focused on RS2Z35 and RS2Z36, belonging to the RS2Z plant-specific clade of SRSF family, whose role in HS response has not been described so far. RS2Z proteins possess an N-terminal RNA recognition motif (RRM) followed by two zinc knuckles (ZnK), a C-terminal region rich in arginine-serine (RS) dipeptides and a serine-proline (SP)-rich region which shows the lowest sequence homology between the two proteins (Fig. 2d; Supplementary Fig. S2) ^23^. *RS2Z35* is a constitutively expressed gene, while *RS2Z36* was identified as DE/DAS gene and is HS-induced in an HSF-dependent manner ^24^, indicating a potential feedback regulation between HSF and SRSFs.

A splicing minigene reporter assay was used to corroborate that the effect of the two RS2Z proteins on *HSFA2* intron 2 was independent of transcription and of splicing of intron 1. The minigene contained the coding sequence of GFP fused to part of the *HSFA2* gene comprised of intron 2 and the flanking exons (Fig. 2b). The minigene reporter was under the control of the HS-inducible HSP21.5-ER promoter ^25^. Co-expression of RS2Z proteins with the minigene reporter resulted in a stronger intron retention compared to mock, both in heat stressed and control protoplasts (Fig. 2d). To investigate the relevance of each domain for the function of RS2Z proteins, C-terminally GFP- or HA-tagged deletion mutants of the RRM, ZnK and RS-rich region were generated (Fig. 2d; Supplementary Fig. S3). Similar to the wild type protein tagged to GFP, all mutants are localized in the nucleus of protoplasts (Supplementary Fig. S4). Based on the minigene reporter assay using the HA-tagged RS2Z variants, deletion of the ZnKs abolished the negative effect of the two proteins on splicing of the *HSFA2* minigene, suggesting that these domains are essential for the function of the proteins as splicing silencers on *HSFA2* (Fig. 2d).

Expression of GFP-tagged RS2Z proteins in tomato transgenic plants under the control of a CaMV35S promoter resulted in the accumulation of the proteins, which showed a nuclear localization in leaf epidermal cells (Supplementary Fig. S5). The endogenous RS2Z35 transcript levels were reduced in both GFP-RS2Z35 and GFP-RS2Z36 transgenic lines compared to WT, while the *RS2Z36* remained unaffected (Fig. 2e). These results indicate an auto- and cross-regulation of *RS2Z35*. Importantly, splicing analysis revealed an enhanced IR for *HSFA2* in both transgenic lines compared to WT after 1 hour of HS (Fig. 2f). The association of *HSFA2* mRNA with both RS2Z proteins in the transgenic lines was confirmed by RNA immunoprecipitation (RIP-qPCR) performed with a nanobody against GFP following formaldehyde cross-linking of RNA-protein complexes in heat stressed-leaves of *CaMV35S::GFP-RS2Z* and *CaMV35S::GFP* transgenic lines (Fig. 2g). In contrast, the RS2Z proteins did not show an enrichment with mRNA from the housekeeping gene *EF1a* which was used as negative control (Fig. 2g).

### RS2Z proteins are important for basal and acquired thermotolerance

To further examine the role of RS2Z proteins in the regulation of splicing and their relevance for thermotolerance, we created single homozygous *rs2z35* and *rs2z36* mutants by CRISPR-Cas9 gene editing. By using two gRNAs for each gene, mutations with either deletion or insertion of one nucleotide were generated (Fig. 3a). This consequently introduced premature termination codons (PTC) through frame shifting (Fig. 3a; Supplementary Fig. S6a). Reduced transcript levels of both *RS2Z* genes in the mutant lines compared to WT indicate degradation by nonsense-mediated mRNA decay (NMD) (Supplementary Fig. S6b). Interestingly, *RS2Z35* levels were increased in the *rs2z36* mutant, while *RS2Z36* levels in the *rs2z35* mutant were similar to WT (Supplementary Fig. S6b).

**Figure 3.**
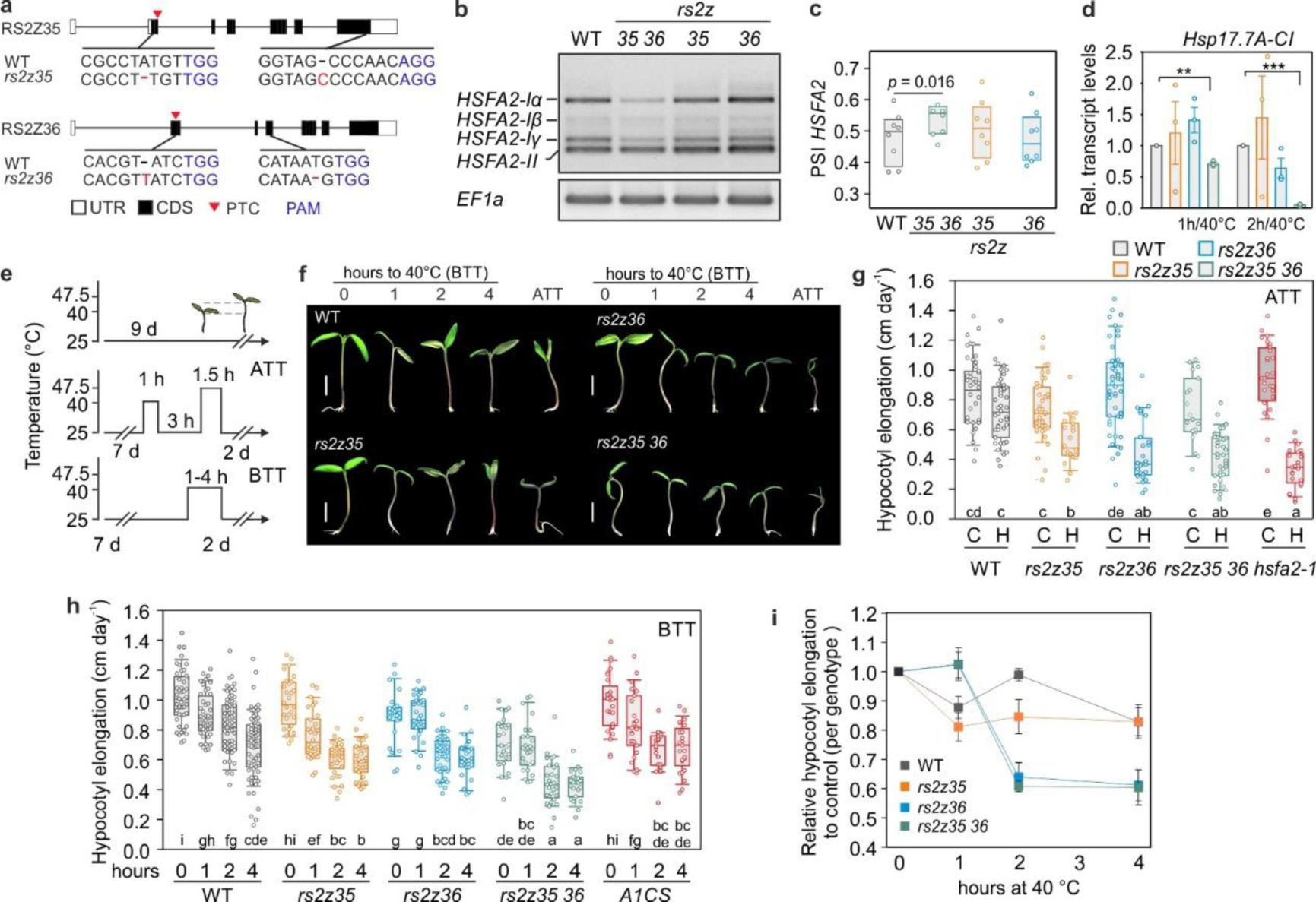
RS2Z proteins are involved in thermotolerance of tomato seedlings. (a) Sequence of *RS2Z35* and *RS2Z36* mutants. Boxes and lines indicate exons and lines introns, respectively. Black regions indicate the coding sequence (CDS) and white untranslated regions (UTR). Red arrowheads depict premature termination codons (PTC) and in blue nucleotides the PAM sequence. (b) Splicing profile of *HSFA2* in WT and single and double *rs2z* CRISPR mutants. *EF1a* is housekeeping gene. The gel is one representative from eight biological replicates. (c) Box plots depict the median, 25% and 75% quartiles of the splicing index (PSI) of *HSFA2* from 8 biological replicates. Statistically significant difference (*p* < 0.05) is shown between the indicated samples. (d) Transcript levels of *Hsp17.7A-CI* based on qRT-PCR analysis. Values are the average of 3 biological replicated, ±SEM. Statistically significant difference based on pairwise *t*-test between the WT and the indicated mutant is depicted by the *p*-value. (e) Heat stress regimes used in thermotolerance assays. BTT: basal thermotolerance; ATT: acquired thermotolerance. (f) Representative images of shoots of WT and mutant tomato seedlings, based on thermotolerance assays indicated on top. Displayed is a compilation from different images chosen to represent average observations. Bar is 1 cm. (g) ATT and (h) BTT of young tomato seedlings from WT and the indicated CRISPR and transgenic knock-down *rs2z* and *hsf* lines. Box plots show the median, 25 and 75% quartiles (*n* = 19–55 for ATT; *n* = 22–79 for BTT) from at least 3 independent experiments. Statistical analysis is based on ANOVA, with Holm-Sidak test for multiple comparisons. Significant differences are indicated by different letters (*p* < 0.05). (h) Relative hypocotyl elongation for each genotype compared to the control (no stress, 0 hour). Each point is the average of 3-7 independent experiments normalized to the control sample of the respective experiment, error bars are ±SEM. For c, g and h, the median is indicated in the box, with the lower upper quartiles, margined by the two extreme values within the interval of 1.5x the interquartile range from the box (whiskers).

The splicing of *HSFA2* was not affected in the single *rs2z35* or *rs2z36* mutants, but was enhanced in a double homozygous mutant line (*rs2z35 36*) obtained by crossing the two single mutants (Fig. 3b,c). This result shows that the two proteins act redundantly as repressors of *HSFA2* intron splicing. In addition, after 1 and 2 hours of HS at 40°C, the *rs2z35 36* mutant exhibited reduced levels of the HSFA2-I-dependent *Hsp17.7A-CI* gene, further linking the function of RS2Z proteins and splicing to the regulation of HSF-dependent gene expression (Fig. 3d) ^9,16^.

Under standard cultivation conditions for tomato, the *rs2z35* and *rs2z36* single mutants were phenotypically identical to the WT plants (Supplementary Fig. S7). Only *rs2z35 36* exhibited a slightly reduced hypocotyl elongation compared to WT seedlings (Fig. 3g,h). The relevance of the two factors for ATT, the capacity of an organism to acclimate and survive an otherwise lethal stress treatment, was examined. Seedlings were pre-acclimated at 40°C for 1 hour, allowed to recover for 3 hours and were then exposed to a challenging HS at 47.5°C for 1.5 hours (Fig. 3e). We used hypocotyl elongation for 2 days following the stress as an indicator of thermotolerance ^9^ and a condition that *HSFA2* gene and particularly *HSFA2-I isoform* is important. Single and double *rs2z* mutants had lower ATT compared to WT seedlings, similar to the *hsfa2-1* mutant (Fig. 3e-g). The higher thermosensitivity of *rs2Z35 36* seedlings in response to an ATT regime is in agreement with reduced *HSFA2-I* transcripts ^16^. However, the reduced ATT in the single RS2Z mutants in the absence of an effect on AS of HSFA2 indicates that the proteins modulate thermotolerance in an HSFA2-independent manner as well.

This result prompted the question whether RS2Z proteins are relevant for basal thermotolerance (BTT), the response of plants to an acute HS. Seedlings of WT and *rs2z* single and double mutants were exposed to 40°C for 1, 2 or 4 hours and then allowed to recover to 25°C for 2 days (Fig. 3 e,f,h). In comparison to WT, the hypocotyl elongation of the *rs2z35* mutant was slightly reduced in seedlings exposed to 1 and 2 hours HS, while *rs2z36* and *rs2z35 rs2z36* mutants exhibited reduced hypocotyl elongation in response to 2 and 4 hour HS treatments (Fig. 3h). In comparison, the seedlings of an *HSFA1a* co-suppression line (A1CS), in which HS response is compromised ^4,26^, were not affected by the short 1 hour HS (Fig. 3h,i). Altogether these results show that the constitutively expressed RS2Z35 is important for thermotolerance during the early stages of HS, in an HSF-independent manner, while the HS-induced RS2Z36 is important for thermotolerance in response to prolonged HS treatments. The lack of a thermotolerance phenotype in 1 hour / 40°C treated *rs2z35 36* seedlings suggests the presence of a compensatory thermotolerance mechanism that is manifested when both genes are mutated.

### RS2Z proteins are core regulators of HS-sensitive alternative splicing

To identify the AS events regulated by the two RS2Z proteins, we performed a global splicing analysis following RNA-Seq on leaves from single and double mutants exposed to 40°C for 1 hour or kept at 25°C as control, and compared them to WT (Supplementary Dataset 4). The comparison in non-stressed plants yielded 170, 174 and 490 bDAS events (IR, ES, A5ss, A3ss) in *rs2z35*, *rs2z36* and *rs2z35 36*, respectively. In HS samples, the number of bDAS events increased to 397, 453 and 729 in *rs2z35*, *rs2z36* and *rs2z35 36* mutants (Fig. 4a). Therefore, the two RS2Z proteins regulate splicing of many genes, both in a specific and redundant manner, accounting for a total of 663 bDAS events under control conditions and 1307 bDAS events under HS (Fig. 4b). Interestingly, there is an overlap of only 126 RS2Z-dependent bDAS events between control and HS, suggesting that they regulate splicing in a temperature-dependent manner. Altogether, the two RS2Z proteins regulate 720 out of 2801 HS-sensitive bDAS genes (Fig. 4c) identified in our initial analysis in WT plants (Fig. 1a). Furthermore, 436 additional bDAS genes, that were not identified as DAS in the WT, are misregulated in at least one of the mutants under HS (Fig. 4c). All these results suggest that RS2Z proteins are core regulators of RNA splicing under HS conditions.

**Figure 4.**
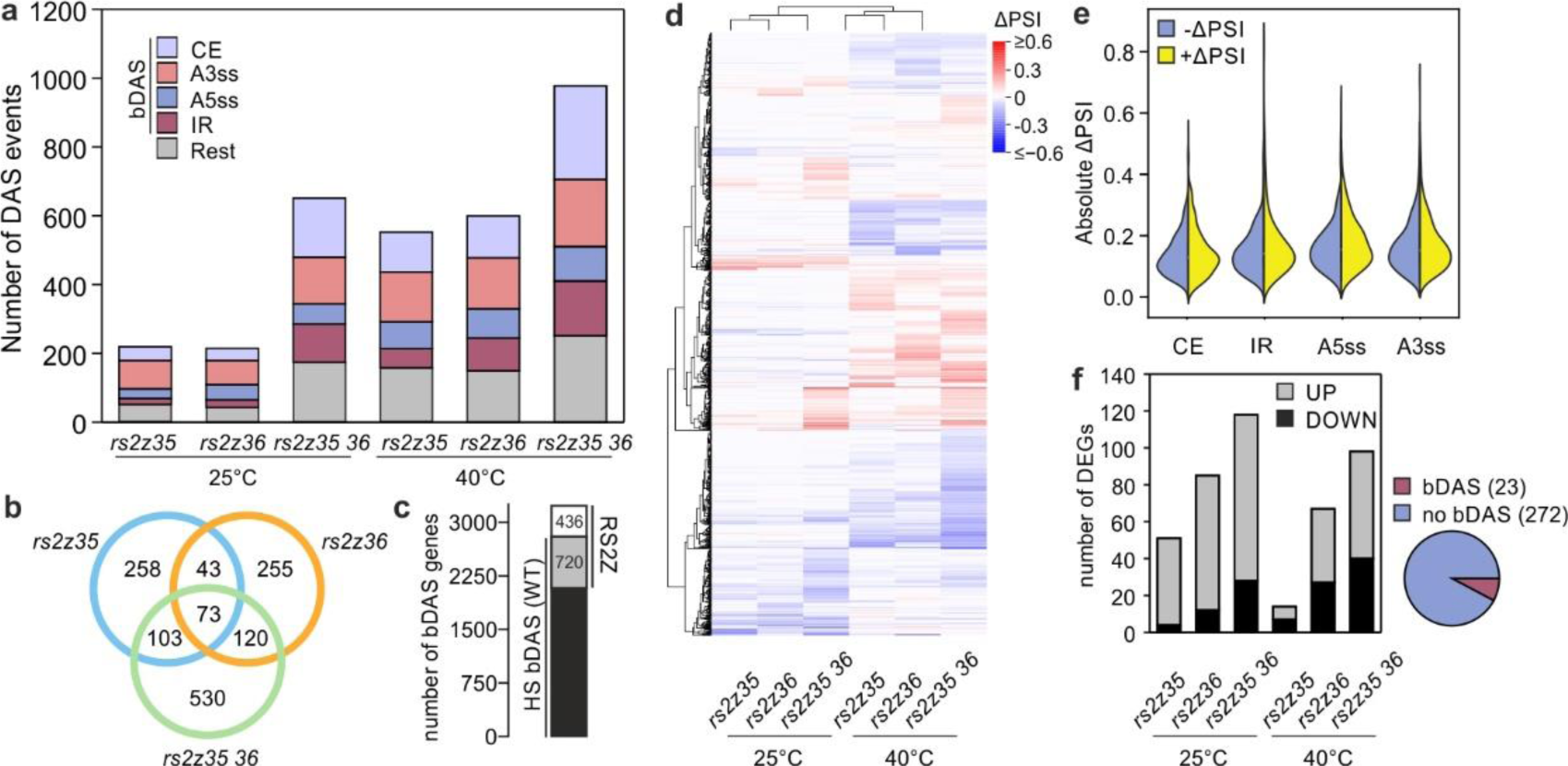
Mutations in RS2Z-coding genes disturb RNA splicing homeostasis. (a) Number of DAS events, categorized based on the splicing type. (b) Overlap and unique DAS events in heat-stressed leaves affected in *rs2z* single and double mutants compared to WT. (c) Overlap of HS-sensitive DAS genes in WT leaves and RS2Z-regulated bDAS genes under HS. (d) Heat map depicting the ΔPSI values of all bDAS events. (e) Distribution of positive and negative ΔPSI values of four binary DAS events in HS leaves of all mutants. (f) Number of DEGs that are up- or downregulated in the single or double mutant lines compared to WT under control or HS conditions. The pie chart depicts the percentage of all DEGs that were identified as DAS based on the comparison of all mutants to WT at any condition.

Based on the splicing quantifications for the bDAS events (relative splice junction usage, delta percent selected index, ΔPSI), we observed that the mutants cause a comparable changes of splicing in both directions (Fig. 4d,e). In addition, the mutants do not show any enrichment of a specific type of alternative splicing. These results indicate that the two proteins are involved in both promoting and silencing splicing and are not preferentially involved in a specific type of splicing but rather contribute to splicing homeostasis. In contrast to the large number of bDAS genes, differential expression analysis between the mutants and WT for each condition yielded a relatively low number of DEGs (Fig. 4f; Supplementary Dataset 5). The majority of the identified DEGs are upregulated in the mutants under control conditions, while under HS the number of induced and reduced genes is similar. Only 23 out of 272 DEGs were bDAS genes indicating that alterations in gene expression are largely independent from splicing.

### The RNA interactome of RS2Z proteins in heat-stressed tomato leaves

Individual-nucleotide resolution UV cross-linking and immunoprecipitation (iCLIP2) ^27^ was employed to generate transcriptome-wide binding maps of the two RS2Z proteins. Young leaves from adult plants of the *PCaMV35S::GFP-RS2Z* transgenic lines were used, along with *PCaMV35S::GFP* as negative control as previously used for RIP-qPCR (Fig. 2g). The leaves were exposed to 40°C for 1 hour and protein-RNA complexes were cross-linked by UV irradiation directly after. Three independent biological replicates were prepared. Immunoprecipitation of cross-linked RNA was done with GFP Trap magnetic beads. iCLIP2 in *PCaMV35S::GFP* plants yielded very low amount of RNA, which was not sufficient for further analysis by RNA-Seq.

iCLIP2 identified a total of 79,257 binding sites (7 nt) for RS2Z35 and RS2Z36, with 49,856 corresponding to RS2Z35 and 53,293 to RS2Z36, and the proteins sharing approximately 50% of their binding sites and roughly 30% of the total binding sites (Fig. 5a; Supplementary Dataset 6). This analysis identified 45 and 33 binding sites for RS2Z35 and RS2Z36, respectively, in both the intronic and exonic regions of *HSFA2* RNA (Fig. 5b). In addition, we identified several binding sites of RS2Z35 on *RS2Z36* mRNA, and RS2Z36 on *RS2Z35* mRNA (Supplementary Fig. S8), in line with cross-regulation of the two proteins. On the global scale the majority of the binding sites are located in coding regions (approximately 67–68%), followed by introns, 5′ and 3′UTRs (Fig. 5c; Supplementary Dataset 6). Similar to *HSFA2*, 7% of the RS2Z-bound RNAs have more than 30 binding sites, while more than 70% of the bound RNAs have up to 5 binding sites (Fig. 5d; Supplementary Dataset 6).

**Figure 5.**
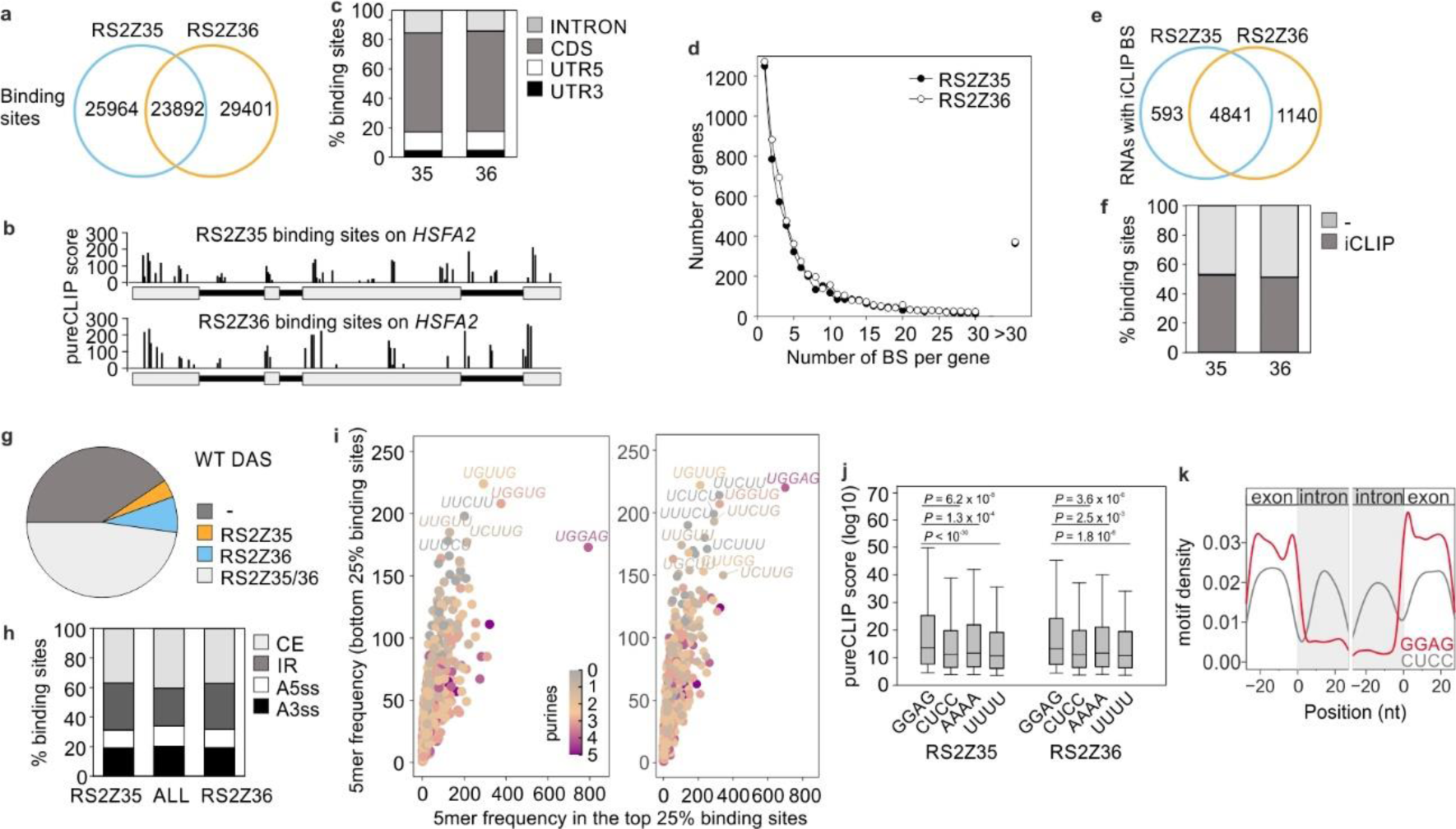
The RNA interactome of RS2Z35 and RS2Z56. (a) Common and specific RS2Z35 and RS2Z36 binding sites identified by iCLIP in leaves of transgenic lines exposed to 40°C for 1 hour. (b) Binding sites and strength shown by PureCLIP score of the two proteins on *HSFA2* mRNA. (c) Fraction of RS2Z35 (35) and RS2Z36 (36) binding sites in introns, CDS or UTRs. (d) Frequency of RS2Z binding sites (BS) on individual genes. (e) Number of genes with iCLIP binding sites (BS). (f) Fraction of putative RS2Z35 (*rs2z35* and *rs2z35 36* DAS) and RS2Z36-dependent (*rs2z36* and *rs2z35 36* DAS) DAS genes identified by iCLIP2. (g) Fraction of WT DAS genes that were identified in RS2Z35 and/or RS2Z36 iCLIP2 analysis. (h) Distribution of alternative splicing events among all HS-sensitive WT DAS genes (ALL) compared to the susets identified as RS2Z35 and RS2Z36 iCLIP2 targets, respectively. (i) Comparison of the frequency of pentamers within the top and bottom 25% of binding sites (left: RS2Z35 binding sites; right: RS2Z36 binding sites). The categorization is based on the binding site strength, defined by the PureCLIP score. (j) PureCLIP score distribution for binding sites containing the GGAG motif within a 15-nt proximity to the binding site centre, in comparison with binding sites featuring control motifs CUCC, AAAA and UUUU within the same window. (k) Positional map of the density of the GGAG (red) and CUCC (grey) motifs in a 51-nt window around the 5′ (left) and 3′ ends (right) of introns.

In total, RS2Z35 bound to 5434 and RS2Z36 to 5981 RNAs, with 73.6% of them being shared between both proteins (Fig. 5e; Supplementary Dataset 7). More than 50% of the putative RS2Z35- or RS2Z36-responsive bDAS genes are bound by these proteins, identified by iCLIP2, suggesting that they directly involved in the regulation of splicing of these RNAs (Fig. 5f). In addition, more than 50% of the HS-sensitive WT bDAS genes are also bound by RS2Z35 and/or RS2Z36, further supporting their central role in alternative splicing during HS (Fig. 5g). The distribution of alternative splicing events among HS-sensitive RS2Z35 and RS2Z36 iCLIP targets was compared with the distribution of AS events in all HS-sensitive WT bDAS genes. We noted a higher enrichment of IR events compared to CE events among RS2Z35 and RS2Z36 iCLIP targets, in comparison to all HS-sensitive WT bDAS genes (Fig. 5h).

To identify the preferential binding motifs of the two proteins, a *de novo* motif discovery analysis was performed using a 51-nt window around the RS2Z binding sites. Motif discovery analysis using XSTREME identified several binding motifs, suggesting that the RS2Z proteins show an ambiguous RNA binding behaviour (Supplementary Dataset 8). To gain further insights into the binding preferences of RS2Z35 and RS2Z36, we examined the sequence composition of the top-25% and bottom-strength 25% binding sites (taking the PureCLIP score as a proxy for binding site strength). For both proteins, the motif UGGAG exhibits a higher frequency in the top 25% binding sites (Fig. 5i). The UGGAG motif is specifically found within the 7-nt binding sites (Fig. 5i), but is not apparent in 17-nt region upstream or downstream of the binding site (Supplementary Fig. S9). This motif is represented in many of the motifs identified by *de novo* discovery (Supplementary Dataset 8) and is similar to purine-rich motifs bound by human SR proteins ^28^.

Based on the de novo motif discovery the GGAG is prevalent while the adjacent upstream nucleotide is variable (Supplementary Dataset 8). The GGAG motif is present in approximately 15% of RS2Z35 and RS2Z36 binding sites and these show significantly stronger binding than binding sites with CUCC (antisense), UUUU or AAAA sequences used as controls (Fig. 5j; Supplementary Dataset 9). Moreover, the presence of two or three GAGG motifs within a 15 nt window results in increased binding strength, something that is not apparent for the CUCC, UUUU or AAAA motifs (Supplementary Fig. S10). Positional analysis of the GGAG motif within a 51-nt window centred on the 5′ and 3′ ends of introns showed that the GGAG motif is predominantly found in the exon region, particularly concentrated right before the 5′ and 3′ of introns (Fig. 5k). Altogether our data suggest that both proteins bind to purine-rich motifs in exons and regulate the splicing of the neighboring intron in a positive or negative manner.

## Discussion

RNA splicing homeostasis is the state of balanced RNA processing where both canonical and alternative splicing pathways operate efficiently and adaptively. This balance is crucial for the accurate and regulated expression of genes under both optimal and stress conditions. Under stressful conditions, alternative splicing confers to the transcriptome landscape and therefore contributes to stress resilience ^29^. Our transcriptome analysis revealed that an exposure to a mild HS treatment results in drastic changes in the splicing profile of many genes (Fig. 1a). Both alternative and canonical splicing are altered to a similar extent (Fig. 1b). We conclude that upon HS, in addition to the well-documented changes in transcript abundance that include both the induction and reduction of many genes, splicing as well is heavily affected in terms of number of RNAs (Fig. 1a,c).

Transcriptional regulation and changes in alternative splicing only partially overlap, as approximately 15% of HS-induced genes are DAS, and 21.5% of DAS are differentially expressed in response to HS (Fig. 1c). Similar observations have been reported in maize^30^, indicating that alternative splicing is an important integral part of the HS response that adds another layer of regulation. DE and DAS genes code for proteins with distinct functions (Fig. 1d). DEGs mainly aid the synthesis of proteins related to stress response, chaperone synthesis and protein folding, while DAS genes code for proteins that have broader roles in the cellular and physiological adjustment that occur during HS. We assume that while DEGs play a direct role in immediate stress response and protection, DAS genes contribute to adaptation processes that enable plants to cope with and recover from HS over time. Worth noticing, proteins involved in pre-mRNA splicing are enriched among DAS genes, which is in agreement with many studies showing that regulators of RNA splicing undergo feedback regulation ^31^. Altogether, the transcriptome analysis suggests that the regulation of alternative splicing is an active process, involving both silencers and enhancers to allow the fine tuning of transcriptome diversity and abundance.

Based on a transient expression system we identified that five SR proteins are putative intron splicing silencers of *HSFA2*, an essential factor for thermotolerance (Fig. 2c) ^16^. We focused on the two proteins that make up the plant-specific RS2Z clade. A tobacco RS2Z gene is involved in plant immunity ^32^, while in rice mutation in *OsRS2Z38* results in changes in alternative splicing under cold stress ^33^. Expression of RS2Z35 and RS2Z36, either transiently in protoplasts or in transgenic plants, results in splicing inhibition of *HSFA2* (Fig. 2c,f). Only the double *rs2z35 36* but not the single mutants show enhanced intron splicing (Fig. 3b), suggesting that they act as splicing silencers of intron 2 of *HSFA2* in a redundant manner. The two proteins share high protein sequence homology in the RRM and ZnK regions. RS2Z proteins are distinct from the other members of the SR protein family due to the presence of the two ZnKs ^23^, which are apparently essential for the function of the two proteins as splicing silencers, likely contributing to RNA binding (Fig. 2d). A previous study in Arabidopsis showed that AtRS2Z33 interacts with other SRSFs and plays a putative role in spliceosome assembly ^34^. We assume that the other SRSFs that affect *HSFA2* splicing can also be part of or affect the activity of the spliceosome complex.

iCLIP2 analysis revealed nearly 50.000 binding sites for each protein with an approximate overlap of 30% (Fig. 5a). This overlap in binding sites is markedly lower than the 75% overlap of bound RNAs, suggesting that the two RS2Z proteins bind to different sites on the same RNAs. Similar to human SRSF proteins ^35^, RS2Z35 and RS2Z36 bind to both exonic and intronic regions (Fig. 5c). The binding of the proteins to RNAs is highly variable in terms of the number of sites per RNA and sequence diversity (Fig. 5d). More than 50% of the HS-sensitive DAS genes identified in WT plants are identified by iCLIP and 25% of them showed a significant alteration in splicing profile in at least one of the mutant lines under HS (Fig. 5e-g). These results demonstrate that the two RS2Z proteins have a central role in regulation of HS-sensitive alternative splicing.

A high number of genes are DAS in the double but not the single mutants (Fig. 4b). This result indicates a possible functional redundancy between the two proteins, but a cooperation between the two paralogs can be assumed as well. Based on iCLIP2 analysis, both proteins can bind to their own but also the others’ mRNAs, and expression of RS2Z35 in transgenic lines results in the endogenous transcript levels of *RS2Z35* and *RS2Z36* (Fig. 2e). All these results suggest a complex auto- and cross-regulation scheme between the two RS2Z proteins that likely fine-tunes the activity of the factors by regulating their expression.

While the two proteins repressed splicing of intron 2 of *HSFA2* (Fig. 3b), the single and double mutant lines showed both enhanced and reduced splicing in other genes, suggesting that they act both as splicing silencers and enhancers (Fig. 4d,e). We assume that this bidirectional regulatory function is dependent on the target RNA and the binding site. Notably, the function of the two proteins as enhancers or silencers is opposite under control and HS conditions in many cases, suggesting that their activity is dependent on interactions with other RNA binding proteins that are involved in splicing which are yet to be found.

Both proteins bind to exons with higher frequency than introns (Fig. 5c) and the purine-rich motif GGAG is enriched in the 5′ and 3′-ends of exons (Fig. 5i-k). iCLIP2 analysis revealed that the presence of more than 1 motif in the window of 15 nt leads to increased binding (Supplementary Fig. S10). A similar purine-rich motif, presented as GAG-repeats, is enriched in exon ends of alternatively spliced pre-mRNAs in heat-stressed *Physcomitrella patens* ^12^, indicating that the mechanism that controls RNA splicing under high temperatures is evolutionarily conserved and might have played a significant role in the ability of plants to adapt to and survive high temperatures. Purine-rich GAAG repeats serve as binding sites for AtSCL33^36^ and might be a general binding site for many plant SRSFs. Furthermore, the human SRSF7, which contains a single ZnK, also binds to a purine-rich motif ^28^.

We have previously shown that alternative splicing of intron 2 of *HSFA2* is important for the synthesis of the HSFA2-I isoform that is involved in ATT ^16^. The *rs2z35 36* mutant that produces less *HSFA2-I* transcripts shows reduced ATT, similar to an *hsfa2* mutant (Fig. 3h). However, the single *rs2z* mutants are more sensitive as well, although they do not exhibit any alteration in *HSFA2* splicing, suggesting HSFA2-independent regulation of stress response and thermotolerance. This is supported further by the reduced thermotolerance of the single and double mutants in BTT treatments in which HSFA2 plays no significant role (Fig. 4g-i).

RS2Z35 is constitutively expressed while RS2Z36 is HS induced in an HSF-dependent manner, indicating a regulatory feedback mechanism between splicing and transcription ^24^. The *rs2z35* mutant is sensitive to 1 and 2 hours of HS while *rs2z36* to 2 and 4 hours of stress, which in combination with the fact that the former is constitutively expressed and the latter stress-induced suggests differing temporal dynamics during HS response (Fig. 4h,i). RS2Z35 is likely to be involved in the initial, rapid response, while RS2Z36 becomes relevant at longer HS durations, likely contributing to later stages of the stress response, by enabling sustained adaptation processes. The increased sensitivity of *rs2z35* mutants to a mild HS treatment, a condition at which the suppression of the master regulator HSFA1a does not cause any significant effect on thermotolerance (Fig. 4h,i), further highlights the importance of the HS-sensitive regulation of RNA splicing.

In summary, our results on the functional relevance of RS2Z35 and RS2Z36 proteins shed light on the crucial role these proteins play in maintaining the homeostasis of RNA splicing under HS. The distinct temporal sensitivity of these proteins to HS, with RS2Z35 being involved in the early response and RS2Z36 in later stages, underscores their importance in different phases of the stress response. The bidirectional role of these proteins, acting as both silencers and enhancers of splicing, further demonstrates their critical function in RNA splicing regulation. This versatility allows for a more refined response to HS, ensuring that the plant can both rapidly react to immediate stress and adapt to prolonged stress conditions. The conservation of the purine-rich RNA-binding motifs across different plant species emphasizes the putative evolutionary importance of these proteins in plant survival under high-temperature conditions, highlighting a direct link between RNA splicing regulation and the ability of a plant to adapt to and survive environmental stresses. We conclude that the HS response and RNA splicing overlap but also work independently to orchestrate the transcriptome landscape to establish thermotolerance as exemplified by the activity of HSFs and RS2Z proteins.

## Methods

### Plant material and stress treatments

All experiments were conducted using *Solanum lycopersicum* cv. Moneymaker as wild type or mutant background. Mutant HSFA2 (*hsfa2-1*) and transgenic HSFA1a lines (HSFA1a co-suppression, A1CS) have been described previously ^4,16^. Plants were grown in the greenhouse in soil supplemented with perlite, with a diurnal cycle of 25°C/16 h with 120 µE illumination and 22°C/8 h. Plants for protoplast isolation were grown in climate chambers with a diurnal cycle of 25°C / 16 h with 120 µE illumination and 22°C/8h on half-strength gelrite-solidified Murashige and Skoog (MS) medium ^37^. Tomato mesophyll protoplast isolation and PEG-mediated transformation were described previously ^4^. In brief, 300,000 protoplasts were transformed with a total of 60 μg plasmid DNA (30 μg HA-SR, 9 μg HSFA2-minigene, adjusted to 60 μg with mock plasmid). 4.5 h after transformation, HS treatments were performed in water baths at the indicated temperatures. Excised leaves were placed on wet tissue paper in sealed petri dishes and subjected to heat stress in water baths. Several leaves were pooled for further RNA and/or protein extraction.

### Seedling thermotolerance assay

Thermotolerance of mutant plants was assessed by hypocotyl elongation under different temperature conditions as previously described ^9^. In brief, seeds of the indicated genotypes were surface sterilized, germinated under dark conditions and subsequently grown on plates containing half-strength Phyto agar-solidified MS medium ^37^ under 25°C/16 h and 22°C/8 h diurnal cycle until approximately 1 cm hypocotyl length. Seedlings were exposed to the indicated temperature regimes and durations. The seedlings were imaged 2 days after the stress treatment, followed by determination of hypocotyl lengths using ImageJ ^38^. The experiment was repeated at least 3 independent times, yielding several independent biological replicates, each one consisting of 5-10 seedlings per treatment. Statistical analysis was performed using one-way ANOVA with Holm-Sidak test for multiple comparisons for *p* < 0.05 using SPSS statistics software (IBM).

### Generation of expression constructs

HA-tagged SR protein constructs have been described previously ^24^. The CaMV35S promoter in pRT-PCaMV35S::GFP-HSFA2 minigene ^16^ has been replaced by the HSP21.5 promoter ^25^ via Gibson assembly ^39^. The generation of RS2Z domain deletion mutants was done by PCR using phosphorylated oligonucleotides and the pRT-HA-RS2Z expression plasmids as template ^40^. The HA-tag was replaced with GFP via NcoI/Acc65I sites to generate GFP-RS2Z expression plasmids. Oligonucleotides used for cloning are listed in Supplementary Table S1.

### Generation of transgenic and mutant plants

For the CRISPR/Cas9-mediated mutation of *RS2Z* genes, two sgRNAs targeting each gene were selected (rs2z35-gRNA1: GGTGGCACACGCCTATATGT, rs2z35-gRNA2: GTGGAAGACGGTAGCCCAAC, rs2z36-gRNA1: GACGGACCCGTTCACGTGATC rs2z36-gRNA2: GTGATGGACGCCGCATAATTG) using CRISPR-PLANT ^41^, and assembly was done using the Golden Gate cloning method ^42^. The plasmids used included the pICSL01009:AtU6p (Addgene under #46968), and those from the MoClo Toolkit (Addgene #1000000044) as outlined by Engler et al. ^43^. The final plasmid, pICSL002208, generously provided by Dr. Nicola Patron from the Earlham Institute in the UK, comprised two sgRNAs each regulated by the *Arabidopsis thaliana* U6 snRNA promoter, the *Streptococcus pyogenes* Cas9 gene driven by the CaMV35S promoter, and the kanamycin resistance gene NPTII under the *NOS* promoter for selection. This vector was introduced into the *Agrobacterium tumefaciens* strain GV3101, which was then used to transform cotyledons of *Solanum lycopersicum* cv. Moneymaker, following the method detailed by McCormick et al. ^44^. Genotyping involved sequencing of the targeted CRISPR/Cas9 editing region. Oligonucleotides used for genotyping are listed in Supplementary Table S1. Experiments were conducted using T3 generation homozygous mutants.

Double *rs2z35 36* mutants were created by crossing homozygous, T-DNA-free single mutants. GFP, GFP-RS2Z35 and GFP-RS2Z36 coding sequences were cloned initially in a pRT vector under the CaMV35S promoter. Following, the whole cassette with the 3′UTR was amplified by PCR (pRT-f: CTAGGTCTCTGGAGTAGGCTTTACACTTTATGC, pRT-r: TACGGTCTCTAGCGTACGGTCACAGCTTGTCTG) to add BsaI sites compatible for Golden Gate cloning into the binary vector pICH86966 carrying the NPTII expression cassette for kanamycin-based selection. Agrobacterium-mediated transformation was done as described for CRISPR. Oligonucleotides used for cloning and genotyping are listed in Supplementary Table S1.

### Immunoblot analysis

For total protein extraction, pelleted protoplasts and homogenized leaves were lysed in high salt lysis buffer (20 mM Tris-HCl, pH 7.8; 0.5 M NaCl; 25 mM KCl; 5 mM MgCl_2_; 30 mM EDTA; 0.5% NP40 substitute; 0.2% sarcosyl; 5% sucrose; 5% glycerol; 0.1% ß-mercaptoethanol; supplemented with cOmplete^TM^ Protease Inhibitor Cocktail, EDTA-free (Merck)), cleared by centrifugation and mixed with 4x Laemmli buffer. Protein concentrations were determined via Bradford assay (Bio-Rad). Equal amounts of proteins were separated by sodium dodecylsulfate (SDS)–polyacrylamide gel electrophoresis (PAGE) and transferred onto 0,45 μm nitrocellulose membranes (GE Healthcare) which were then probed with the following primary and secondary antibodies with the indicated dilutions: αHemagglutinin (HA, 1:2000, BioLegend), αGFP (1:5000, Roche), αHSC70 (Spa-820, 1:10,000, StressGen Biotechnologies), αMouse IgG, HRP-conjugated (1:10,000, Sigma-Aldrich), αRabbit IgG, HRP-conjugated (1:10,000, Sigma-Aldrich). HRP signals were obtained using the ECL Kit (Perkin-Elmer Life Sciences) followed by imaging using the ECL ChemoStar 6 instrument (Intas Science Imaging).

### Microscopy analysis

Cells from the adaxial side of leaves from GFP or GFP-RS2Z transgenic plants or protoplasts transformed with GFP-containing constructs were imaged using a confocal laser-scanning microscope (Zeiss CLSM 780, Plan Apochromat 20-64x objectives). 100,000 Protoplasts were transformed with 10 µg GFP-RS2Z plasmid and 10 µg of pRT-2xCaMV35S::ENP1-mCherry (or as control 4 µg pRT-CaMV35S::GFP, 10 µg pRT-2xCaMV35S::ENP1-mCherry, 6 µg mock plasmid). Protoplasts were incubated overnight at 25°C. The excitation and emission spectra of GFP were set as 488 nm and 490–548 nm, for mCherry 561 nm and 570 - 656 nm, and chlorophyll B 633 nm and 665 - 738 nm, respectively. Images were acquired using Zen software (Zeiss).

### GFP nanobody purification

The GFP nanobody used for RIP was obtained through bacterial expression following transformation of *E. coli* BL21 Star (DE3) pRARE (Thermo Fisher Scientific) with Addgene plasmid #49172, encoding for IPTG-inducible expression of a His-tagged GFP nanobody. In brief, nanobodies were expressed in 2 L bacterial culture for 16 h at 37°C post IPTG-induction and supplemented with 1 mM PMSF before harvest. Cell pellets were resolubilised in lysis buffer (50 mM HEPES, pH 7.6; 0.5 M NaCl; 0.1x protease inhibitor cocktail (Roche); 5% glycerol, 2 mM ß-mercaptoethanol), treated with DNaseI and lysed with a French Press cell disruptor (Thermo Fisher Scientific). Nanobodies were purified from the cleared lysate via gravity flow column purification using Ni-NTA agarose (Qiagen) and eluted with imidazole. Purified nanobodies were further subjected to a buffer exchange and were passed through a centrifugal filter (Amicon^®^Ultra-15, Millipore) with a 50 kDa cutoff, allowing nanobodies to pass through while retaining larger *E. coli* protein contaminants.

### RNA isolation, cDNA synthesis and qRT-PCR

Total RNA from protoplasts and homogenised leaves was isolated using TRIzol (Invitrogen) following the manufacturer’s instructions. Up to 1 µg total RNA was treated with DNaseI (Roche) and further used for first-strand cDNA synthesis using the RevertAid Reverse Transcriptase (Thermo Fisher Scientific) and oligo(dT) primers. Reverse transcriptase PCRs (RT-PCR) to visualize splicing patterns were conducted using Pfu polymerase, 10 ng cDNA and up to 28 PCR cycles to avoid saturation. PCR products were subsequently analyzed via agarose gel electrophoresis and the ratio of splice variants was assessed using ImageLab (Bio-Rad) or Fiji/ImageJ. Quantitative real-time PCR (qRT-PCR) was performed in at least technical duplicates and a minimum of 3 biological replicates using the PowerUp SYBR Green Master Mix (Thermo Fisher Scientific) on a StepOne Plus Real-Time PCR system (Thermo Fisher Scientific). Relative transcript levels were determined using the 2-ΔΔCt method ^45^ by normalisation against *EF1α* as internal control as well as against the respective biological control. Oligonucleotides used to amplify transcripts of interest via RT-PCR and qRT-PCR are listed in Supplementary Table S1.

### RNA immunoprecipitation (RIP)

The RIP procedure was done similarly to Köster et al. ^46^ and performed in 4 independent biological replicates. Young leaves from 8-week-old transgenic plants were subjected to 1 h 40°C in sealed petri dishes in a water bath. Following HS, leaves were subjected to formaldehyde (0.5% in PBS) crosslinking for 20 min. Per sample, 1 g crosslinked plant material was homogenized and subsequently lysed by addition of 2 volumes RIP lysis buffer (50 mM Tris-HCl pH 7.6, 4 mM MgCl_2_, 500 mM NaCl, 1% NP40 substitute, 1% sodium deoxycholate, 10% glycerol, 0.1% SDS, 1 mM NaF, 5 mM DTT, 1 mM PMSF) supplemented with Protease inhibitor cocktail (Roche) and 100 U/ml Ribolock (Thermo Fisher Scientific), and brief incubation at 40°C and 1,400 rpm, followed by 5 min incubation on ice with frequent vortexing. Following clearing via centrifugation and filtration, the lysates were diluted by addition of 2 volumes lysis buffer without salt and detergents. An aliquot of the lysate was taken as input sample. For immunoprecipitation (IP), purified His-tagged GFP-nanobody was bound to PureCube Ni-NTA Magbeads (Cube biotech) in the presence of 500 ug/ml Heparin. IP was carried out for 1 h at 4°C with end-over-end rotation. Afterwards, the beads were washed six times with wash buffer (lysis buffer supplemented with Protease inhibitor cocktail (Roche) and 50 U/ml Ribolock (Thermo Fisher Scientific)) and a final time with last wash buffer (50 mM Tris-HCl pH 7.6, 4 mM MgCl_2_, 150 mM NaCl, 1 mM NaF, 5 mM DTT, Protease inhibitor cocktail (Roche) and 50 U/ml Ribolock (Thermo Fisher Scientific)). Finally, proteins were eluted using imidazole-containing elution buffer (25 mM Tris-HCl pH 7.6, 150 mM NaCl, 5 mM EDTA, 400 mM Imidazole, Protease inhibitor cocktail (Roche), 100 U/ml Ribolock (Thermo Fisher Scientific)). RNA from input and elution samples were extracted using TRIZol (Invitrogen) following the manufacturer’s instructions. Reverse crosslinking was achieved by incubation at 65°C for 7 min at 1,400 rpm upon addition of the TRIZol reagent. cDNA synthesis and qRT-PCR were carried out as described earlier. Relative enrichment was calculated using the 2^−ΔΔCt^ method ^45^ by normalizing the elution of each genotype to its corresponding input and subsequent normalization of RS2Z samples to GFP. Undetermined ΔCt values were set to 40 to allow calculation of fold enrichment (observed in the case of *EF1α* transcripts but not *HSFA2*).

### Transcriptome analysis by RNA-Seq

RNA-Seq was performed in 3 biological replicates. Excised leaves from six 6-week-old wild type and mutant plants were exposed to 1 h 40 °C (control: 1 h 25°C) and total RNA was extracted using TriZol (Invitrogen) followed by DirectZol (Zymo Research) column purification including DNaseI digestion. RNA integrity was evaluated using the Qubit IQ Assay kit (Thermo Fisher Scientific) and RNAs with IQ scores > 8.3 were used for library construction with the NEBNext Ultra II non-directional RNA Library Kit with polyA selection, and subsequent paired-end RNA-sequencing with 150 nt each direction on an Illumina NovaSeq 6000 machine. Library construction and sequencing were performed by Admera Health, LLC (New Jersey, USA). Quality control was performed using FastQC. Reads were mapped against the tomato reference genome (*S. lycopersicum* cv. Heinz 1706 genome build SL4.0) with the corresponding ITAG4.0 annotation ^47^ using Star (Galaxy version 2.7.8a) ^48^ with soft clipping and 5% mismatched bases allowed. For DEG analysis, gene expression was quantified as absolute read count for each coding gene using htseq count (Galaxy version 0.9.1) ^49^. Downstream analyses were performed in R (version 4.1.1; ^50^). Batch effects in the heat stress samples were identified by principal component analysis (PCA) and removed using ComBat-Seq (version 3.42.0) ^51,52^. Prior to differential expression analysis, low abundance transcripts (read sum across all samples < 10) were removed. Differential expression analysis was performed using DESeq2 (version 1.32.0) ^53^. log2FC shrinkage was performed using the apeglm package (version 1.14.0) ^54^ to avoid overestimation of fold changes in low expressed genes or those with high dispersion. Transcripts were assigned as significantly differentially expressed when their expression levels relative to the control were |log2FC| > 1 with an FDR-adjusted *p*-value of < 0.01. RNA-Seq datasets are deposited at the NCBI Gene Expression Omnibus (GEO, www.ncbi.nlm.nih.gov/geo/) under the accession number GSE261816 (superseries GSE262552).

### Alternative splicing analysis

The alternative splicing analysis was performed using the MAJIQ and VOILA packages (both version 2.3) ^55^. First, *majiq build* was used to generate a splice graph for each gene and to detect so-called local splicing variations (LSVs) in the splice graphs by taking the BAM files described earlier and the ITAG4.0 corresponding GFF3 file as input. In the second step, changes in the splice junction usage and retention levels of introns (ΔPSI) were quantified between control and HS conditions for the WT and RS2Z single and double mutant datasets using *majiq deltapsi*. In the last step, binary splicing events (e.g. cassette exon or intron retention events) were reconstructed from the previously identified LSVs with *voila modulize*, which also computed the probability that the estimated |ΔPSI| of a LSV exceeds the selected cutoff of 2% (--changing-between-group-dpsi 0.02). Overall, 14 distinct binary events are reported, that have either been reconstructed from a single (e.g. intron retention events) or two LSVs (e.g. cassette exon events), whereby for each LSV two junctions are reported by *voila modulize*. Alternative splicing events were considered significantly regulated between two conditions if: (i) both junctions of at least one LSV have |*P*_(|ΔPSI| ≥ 0.02)_ ≥ 0.9 and |ΔPSI| ≥ 0.05, (ii) all involved junctions have |ΔPSI| ≥ 0.025, (iii) junction pairs of the same LSV have inverse regulation and (iv) the lower |ΔPSI| in a junction pair must be at least 50% of the higher |ΔPSI|. The heat map with the ΔPSI values was prepared using the SRplot platform ^56^.

### individual-nucleotide resolution cross-linking and immunoprecipitation (iCLIP)

iCLIP2 (individual-nucleotide resolution UV cross-linking and immunoprecipitation) was performed in three biological replicates following the procedure described by Buchbender et al. ^27^ with modifications to meet the adaptions made for plants described by Meyer at al. ^57^. Young leaves from 8-week-old transgenic plants were subjected to 1 h 40°C, subsequently irradiated with 750 mJ/cm^2^ UV light at 254 nm (CL-1000, UVP) on ice and immediately snap-frozen in liquid nitrogen. Lysates from 1.2 g plant material each were prepared as described for RIP but using CLIP lysis buffer (50 mM Tris-HCl pH 7.5, 4 mM MgCl_2_, 500 mM NaCl, 0.25% NP40 substitute, 0.25% sodium deoxycholate, 1% SDS, 5 mM DTT, protease inhibitor cocktail (Roche), 1 mM PMSF, 10 mM Ribonucleoside Vanadyl Complex (NEB) and 100 U/ml Ribolock (Thermo Fisher Scientific)). Lysates were cleared and subsequently diluted with 2 volumes CLIP lysis buffer without NaCl, followed by incubation with GFP-Trap magnetic agarose beads (ChromoTek) for 1.5 h at 4°C with end-over-end rotation. The beads were washed four times using plant IP wash buffer (50 mM Tris-HCl pH 7.5, 4 mM MgCl_2_, 500 mM NaCl, 0.5% NP40 substitute, 0.2% sodium deoxycholate, 1% SDS, 2M urea, 2 mM DTT, protease inhibitor cocktail (Roche)) and once using PNK Wash buffer ^27,57^. The beads were then subjected to a DNase treatment using 2 μl Turbo DNase (Invitrogen), as well as an RNase treatment using 5 μl Ambion RNaseI (Invitrogen) in 1:6000 dilution (0.083 U/sample) for 10 min at 37°C, followed by two washes with high salt wash buffer and 3 washes with PNK wash buffer ^27^. Further steps including library preparation were carried out according to Buchbender et al. ^27^. In brief, the RNA was 3’end dephosphorylated and an L3 linker was ligated. Next, the RNA-protein complexes were separated via denaturing gel electrophoresis and subsequently transferred onto a 0,45 µm nitrocellulose membrane. In a pilot run, the 5′ ends were radio-labelled prior to gel electrophoresis to assess the desired region to be excised. The region 20– 60 kDa above the expected size of the respective fusion protein was excised from the membrane and the RNA was liberated by proteinase K treatment, followed by RNA isolation and cDNA synthesis using SuperScript™ III Reverse Transcriptase (Invitrogen). The cDNA was subjected to NaOH treatment and clean-up, followed by second adapter ligation. Next, the cDNA was pre-amplified with 6 cycles using Phusion HF Mastermix (New England Biolabs) and size-selected using the ProNex Size-Selective Purification System (Promega). cDNA libraries were then PCR-amplified with 13 cycles and the final library quality was assessed using a TapeStation System (Agilent Technologies), followed by a second size selection. Purified iCLIP2 libraries were sequenced on a NextSeq500 machine (Illumina) using a NextSeq^®^ 500/550 High Output Kit v2 as 75-nt single-end reads yielding between 4.6 (minus UV controls) and 34.3 million reads. iCLIP2 datasets were deposited at the NCBI Gene Expression Omnibus (GEO, www.ncbi.nlm.nih.gov/geo/) under the accession number GSE260668 (superseries GSE262552).

### iCLIP data analysis

iCLIP2 sequencing data were processed as described in Busch et al. ^58^ with minor changes. In brief, reads with a sequencing quality below 10 in the barcode regions were removed, followed by adapter trimming with an accepted error rate of 0.1 on the right end and a minimal required overlap of 1 bp between reads and adapter. Trimmed reads with a minimal length of 15 bp were kept and subsequently mapped against the tomato reference genome (*S. lycopersicum* cv. Heinz 1706 genome build SL4.0) with the corresponding ITAG4.0 annotation ^47^ using Star (version 2.7.3a) ^48^ without soft clipping on the 5’ ends of the reads and with 4% mismatched bases allowed. Uniquely mapped reads were de-duplicated using UMI-Tools (version 1.1.2) ^59^. The processed reads from all replicates per genotype were merged into a single file and subjected to peak calling using PureCLIP (Galaxy version 1.0.4) ^60^ with default paraments. Downstream analyses were performed in R (version 4.1.1) ^50^. Binding site identification based on crosslink sites was carried out using BindingSiteFinder (version 1.7.9) ^27^. Prior to binding site definition, the crosslink sites were filtered to remove those with the lowest 5% PureCLIP scores. Subsequently, a gene-wise filter was applied to discard sites with the lowest 10% PureCLIP score on each gene, ensuring a fair filtering approach for genes irrespective of gene expression levels.

The remaining sites were merged into 7 nt wide binding sites, excluding those with a width of less than 3 nt and those with fewer than two cross-link events. Maximum PureCLIP scores were re-assigned to binding sites to obtain the respective PureCLIP score for each binding site. All binding sites were controlled for reproducibility, requiring support from all three replicates. A replicate was considered to support a binding site if it harbors more cross-links than the replicate-specific threshold, defined by the 10% quantile of the cross-links distribution and necessitating a minimum 2 crosslinks in all binding sites. Binding sites were overlapped with gene and transcript annotations from ITAG 4.0. Binding sites within protein-coding genes were assigned to the following transcript regions: intron, coding sequence, 3′- or 5′-UTR. Binding sites that overlapped with multiple transcript regions were resolved by selecting transcript region most frequently observed. Any sites overlapping with multiple genes were removed from the analysis.

For genes without UTR annotations, 5′ and 3′UTR regions were included by extending the genes by 200 nt upstream of the start codons and 300 nt downstream of the stop codons, respectively. The extension length for regions was decided based on the average length of 5′ and 3′UTRs in the tomato reference genome.

### Motif analysis

The sequence content around binding sites was analyzed by counting the frequency of pentamers (5-mers) using Biotrings::oligonucleotideFrequency (version 2.70.1) ^61^ within the 7 nt binding site and the flanking 22 nt regions both upstream and downstream of the binding sites. Selection of the top and bottom 25% binding sites was carried out by filtering for those with PureCLIP score above the 75^th^ percentile and below the 25^th^ percentile cutoff, respectively. The *de novo* motif discovery and enrichment analysis were performed using the XSTREME tool from the MEME suite ^62^. To identify potential RBP binding motifs, the binding sites were extended by 25 nt from the binding site center, resulting in a 51 nt window. Sequences within this window were extracted using Biostrings (R package; version 2.70.1) and used as an input for the analysis. Shuffled input sequences served as a control. The XSTREME analysis was carried out with default parameters.

To analyze motif occurrences, the binding sites were extended by 7 nt on either side from the center. The numbers of GGAG motif and the control motifs (CUCC, AAAA and UUUU) within the binding site window (15 nt) were compared with the respective PureCLIP scores. For relative motif position calculation, a 51 nt window centered on the 5′ and 3′ ends of introns was selected. Subsequently, the genomic coordinates of the GGAG and CUCC motifs within the selected window were extracted. The relative positions of the motifs within the regions were then calculated by measuring the distance between the motif start and the 5′ and 3′ ends of the introns.

### Protein sequence alignment

Pairwise sequence alignments of RS2Z35 and RS2Z36 were performed using EMBOSS Needle with default settings (matrix: EBLOSUM62, Gap_penalty: 10.0, Extend_penalty: 0.5), employing the Needleman-Wunsch alignment algorithm ^63^.

### Gene ontology enrichment analysis

Gene ontology was analyzed using PANTHER v14 using the ITAG 4.0 annotation ^64^. The *P*-value for each category was calculated based on Fischer’s exact test and Bonferroni correction for multiple testing.

### Data availability

The RNA-Seq and iCLIP2 data generated in this study have been deposited at the NCBI Gene Expression Omnibus (GEO, www.ncbi.nlm.nih.gov/geo/) under the SuperSeries GSE262552.

## Acknowledgements

We thank Dorothee Staiger for the helpful comments during iCLIP preparation, Holger Schranz for plant cultivation in the greenhouse and Kerstin Zehl for technical assistance in protoplast preparation. We acknowledge the support of DFG with the grant FR 3776/5-1 to S.F., SFB902 to E.S., and BMLS Josef Buchmann PhD Scholarship to R.R.E.R. Support by the IMB Genomics Core Facility and the use of its NextSeq500 (funded by the Deutsche Forschungsgemeinschaft (DFG, German Research Foundation) – 329045328) as well as the help of Anke Busch are gratefully acknowledged.

## Author contributions

S.F. and R.R.E.R.: conceptualization; R.R.E.R, S.V., M. K., S.S., M.B., C.B. and K.Z.: data curation; R.R.E.R, S.V., M. K., S.S., F.M., K.L., D.B., T.B., M.M.M. and K.Z.: formal analysis, investigation and methodology; S.F. and E.S.: funding acquisition; S.F., M.M.M., K.Z., M.F. and E.S.: supervision; S.F., R.R.E.R., S.S., M.K. and S.V.: validation; S.F., R.R.E.R., S.V., S.S. and M.K.: visualization; S.F., R.R.E.R., S.V. and S.S.: writing original draft; R.R.E.R, S.V., M.K., S.S., F.M., K.L., D.B. M.B., M.C., T.B., C.B., M.F., E.S., M.M.M., K.Z. and S.F.: writing—review and editing.

## Additional information

**Supplemental Table S1.**
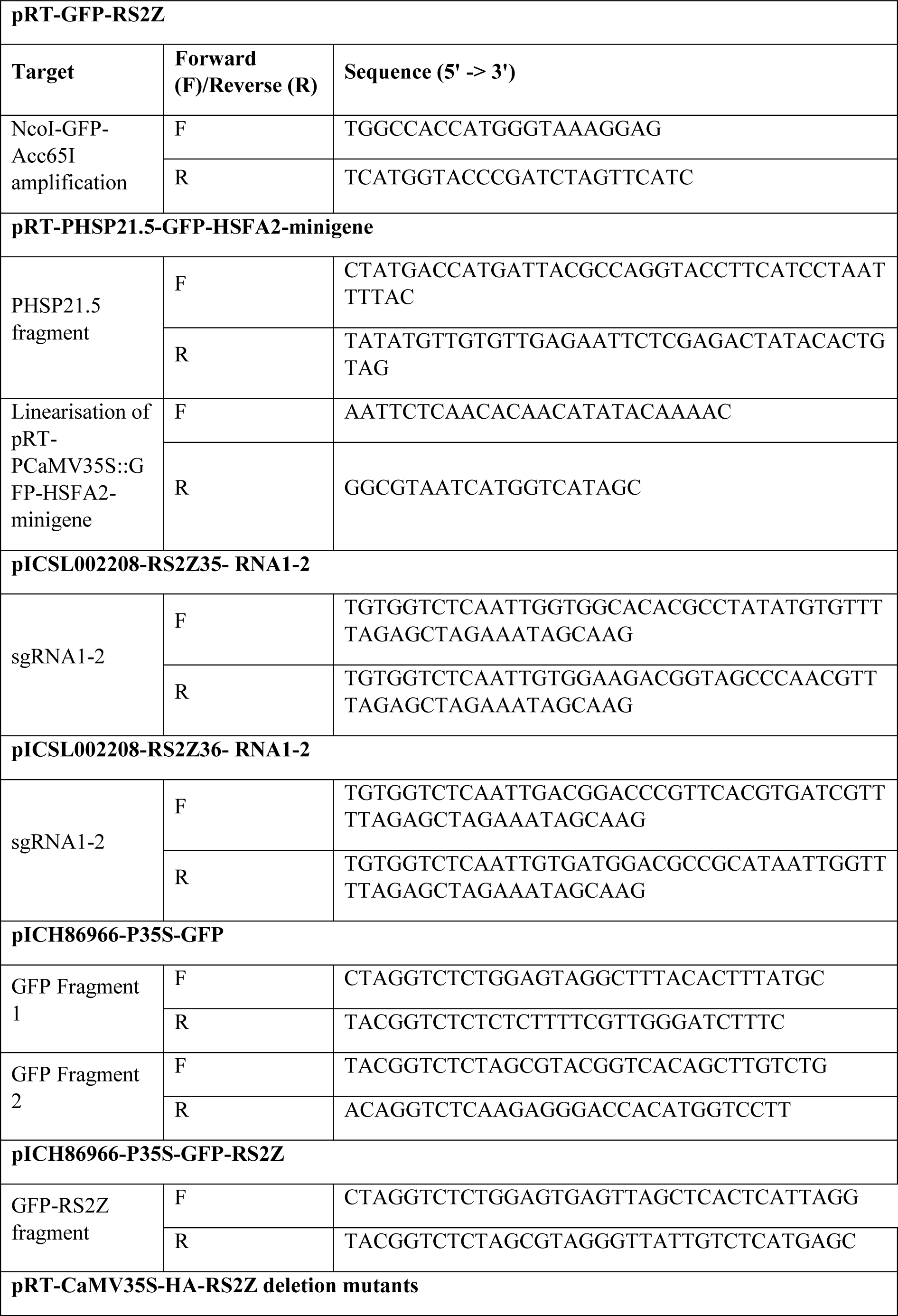
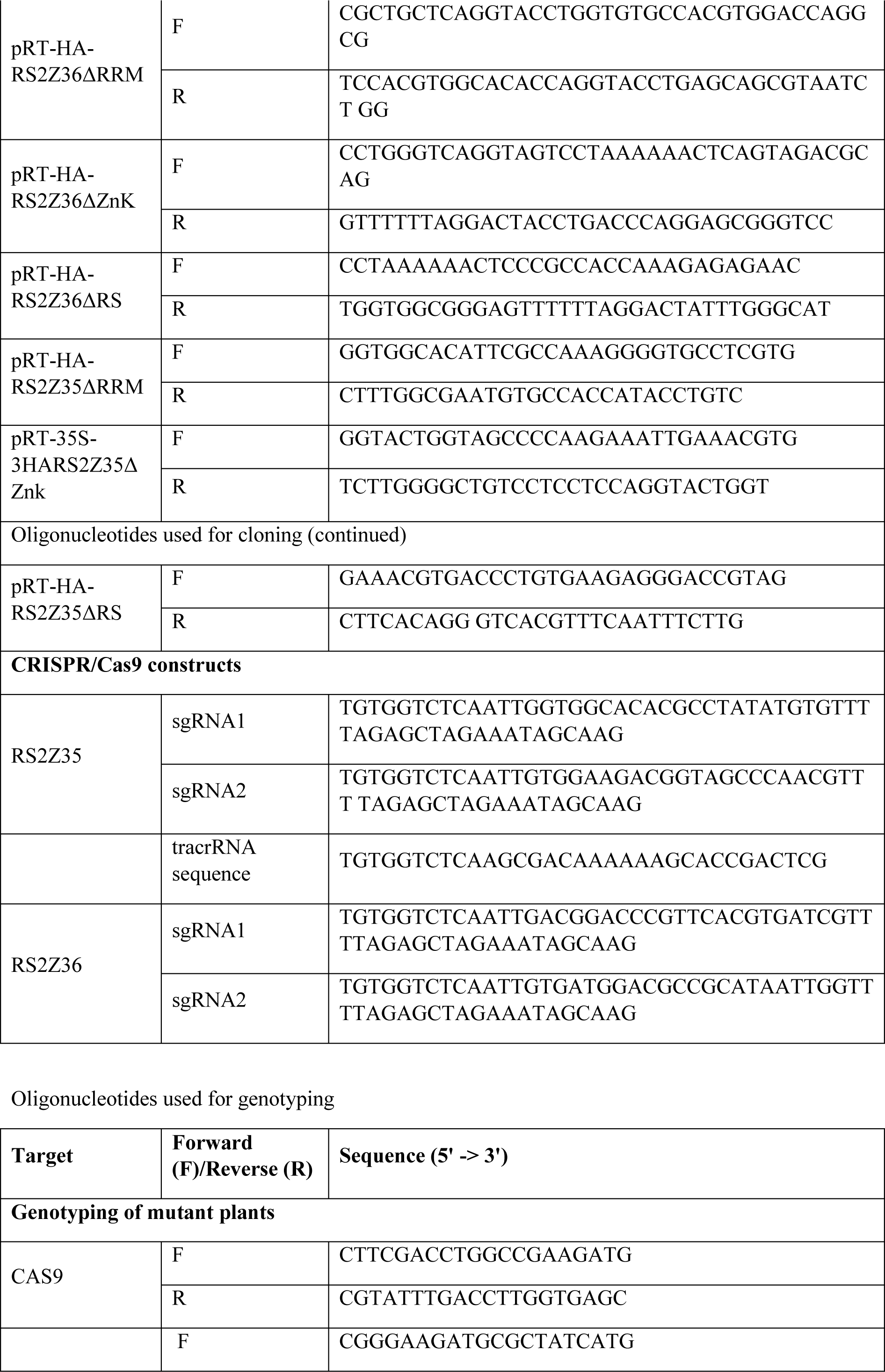
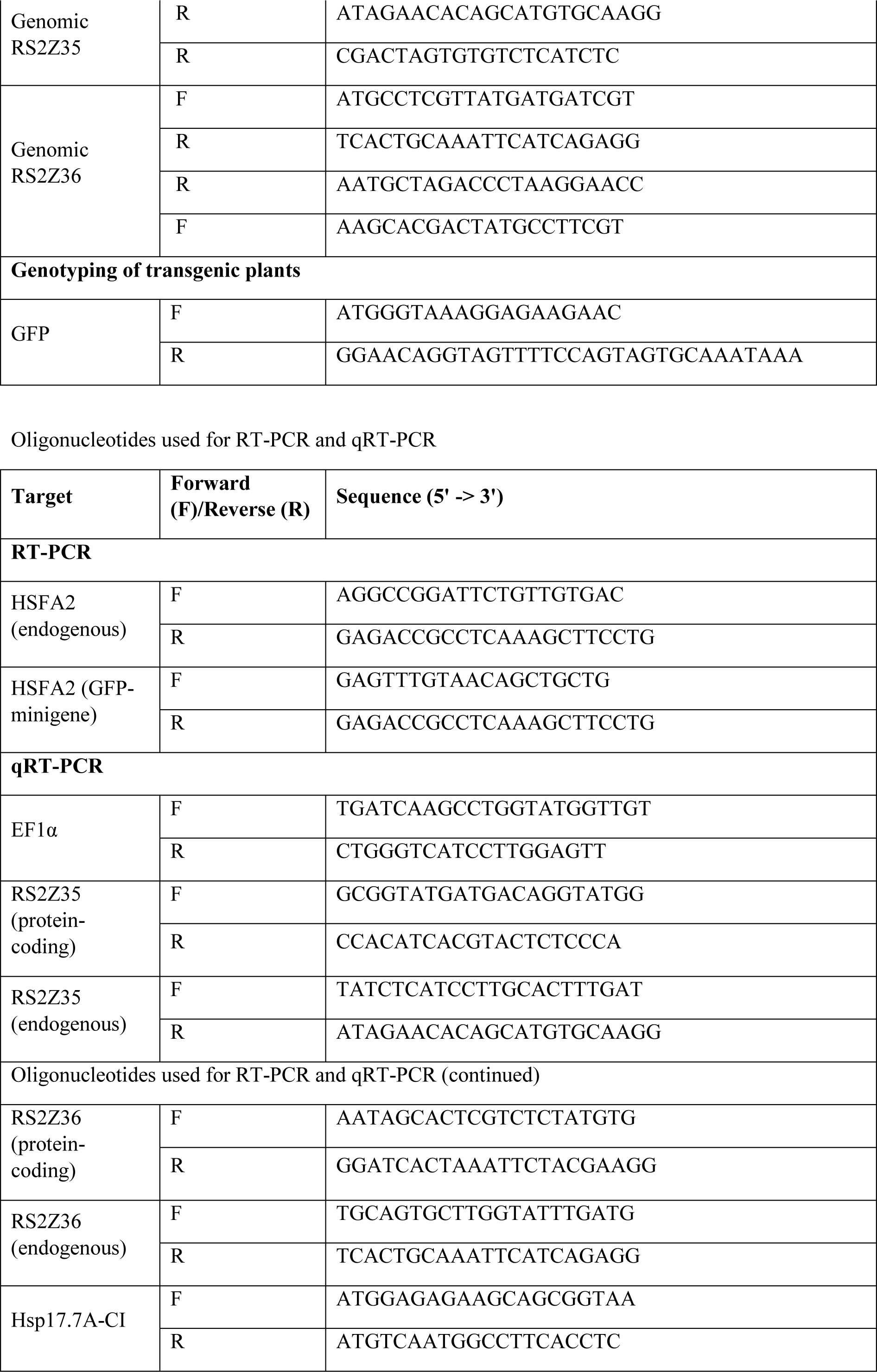
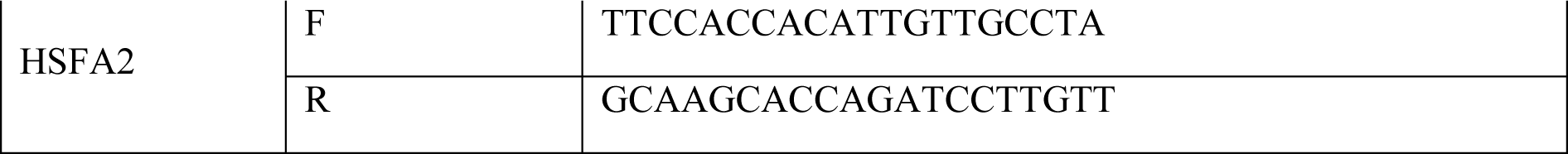
List of oligonucleotides.

## Supplementary Datasets

**Supplementary Dataset S1.** List of alternative splicing events in leaves of wild type plants under control and heat stress conditions.

**Supplementary Dataset S2.** List of differentially expressed genes in leaves of wild type plants under control and heat stress conditions.

**Supplementary Dataset S3.** Gene ontology term of DEG and DAS genes wild type plants under control and heat stress conditions.

**Supplementary Dataset S4.** List of alternative splicing events differentially regulated in in leaves of single and double *rs2z* mutants compared to wild type plants under control and heat stress conditions.

**Supplementary Dataset S5.** Differentially expressed genes in tomato leaves of single and double *rs2z* mutant lines compared to wild type under control and heat stress conditions.

**Supplementary Dataset S6.** Binding sites of RS2Z35 and RS2Z36 on RNA of heat-stressed tomato leaves.

**Supplementary Dataset S7.** List of tomato genes with RS2Z35 and RS2Z36 binding sites

**Supplementary Dataset S8.** List of nucleotide motifs identified by de novo motif search for RS2Z35 and RS2Z36 binding sites.

**Supplementary Dataset S9.** Binding sites of RS2Z35 and RS2Z36 on RNA of heat-stressed tomato leaves and occurrence of GGAG, CUCC, UUUU and AAAA motifs.

## Supplementary Figures

**Supplementary Figure S1.**
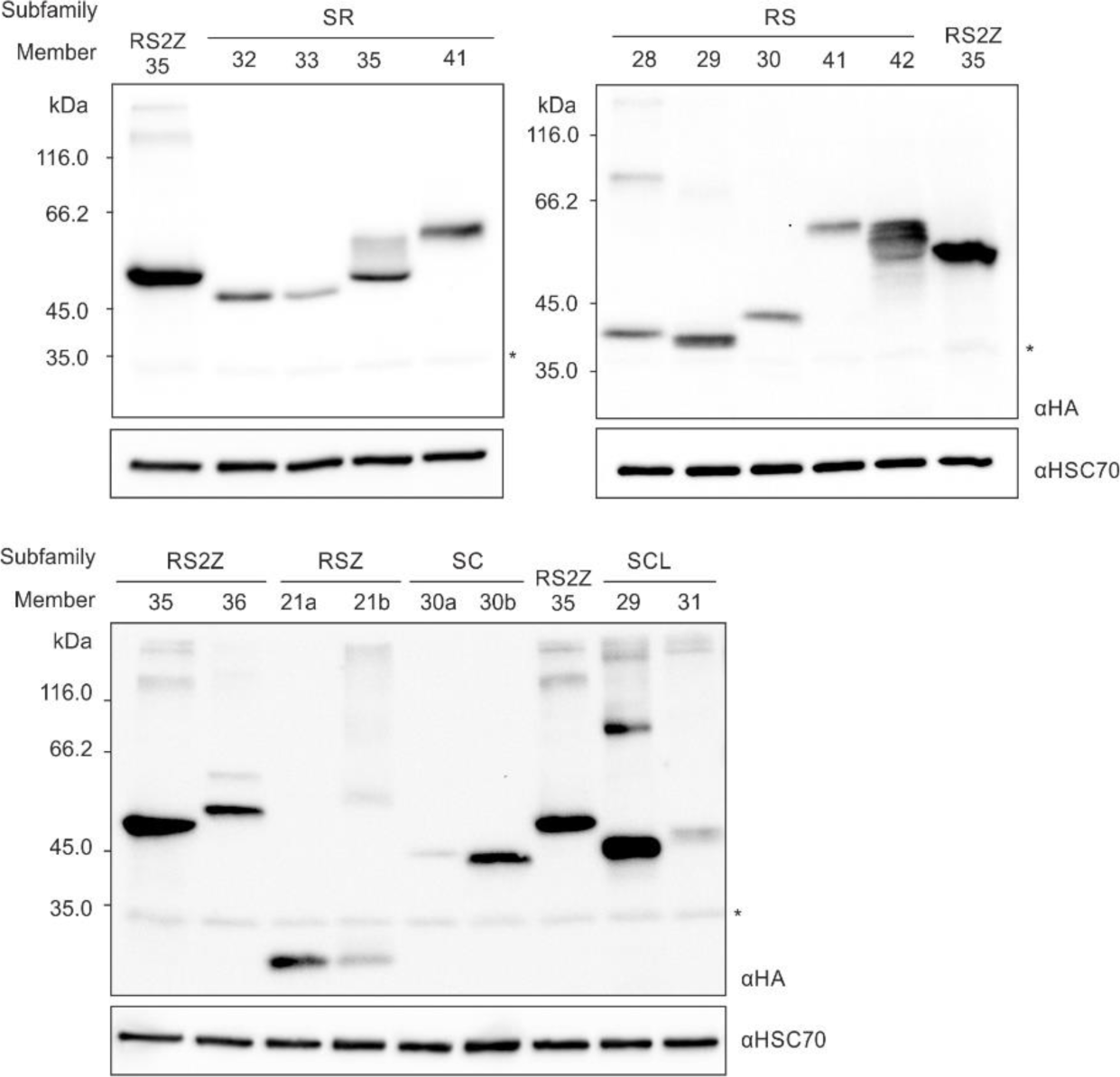
Immunoblot analysis of HA-tagged SR proteins expressed in tomato mesophyll protoplasts. RS2Z35 is shown in all immunoblots for comparison. HSC70 is shown as loading control. The blots are Supplementary to Fig. 2c.

**Supplementary Figure S2.**
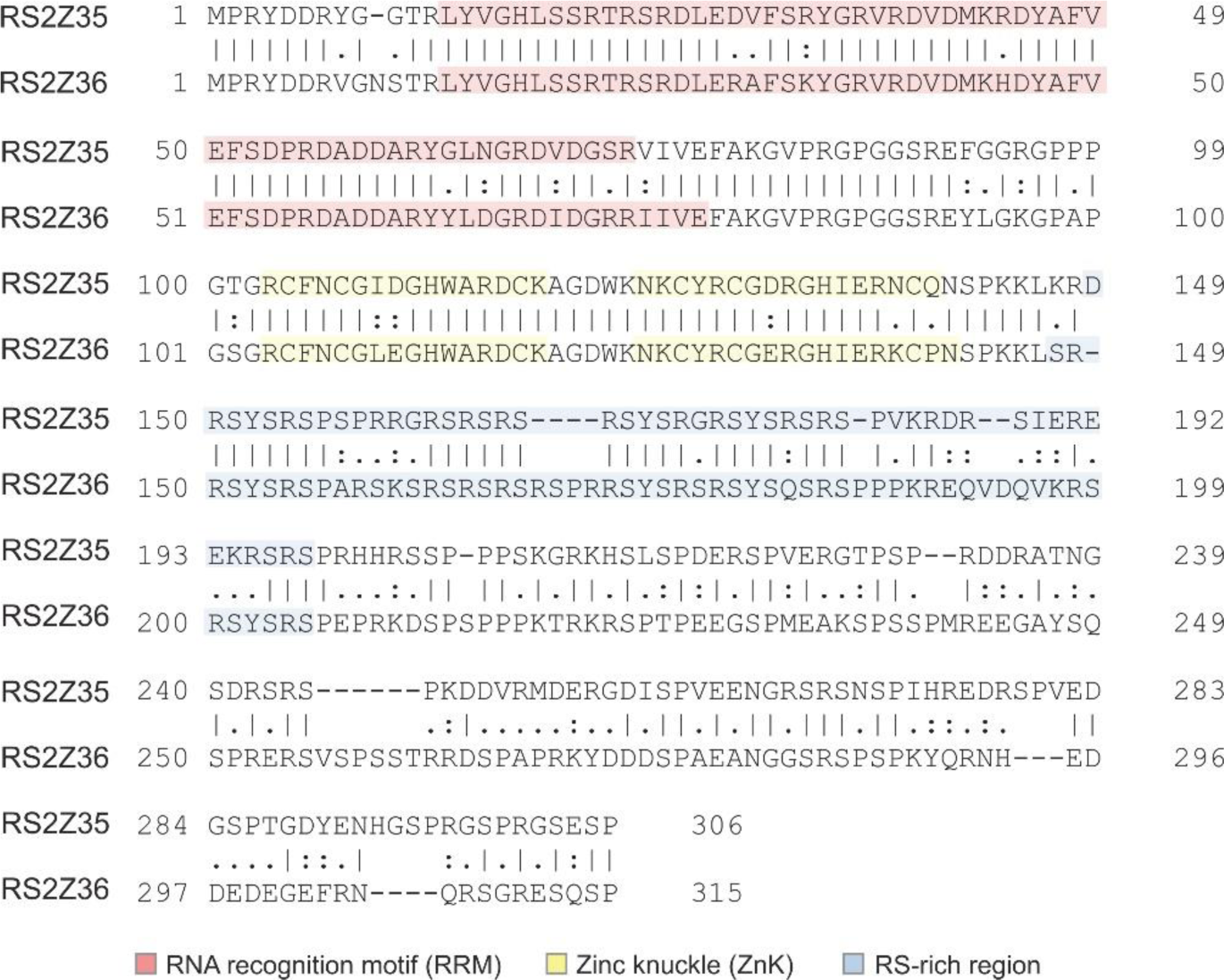
Alignment of the protein sequence of RS2Z35 and RS2Z36. Red color indicates the RRM, yellow the ZnKs and blue the RS-rich region. This figure is Supplementary to Fig. 2d.

**Supplementary Figure S3.**
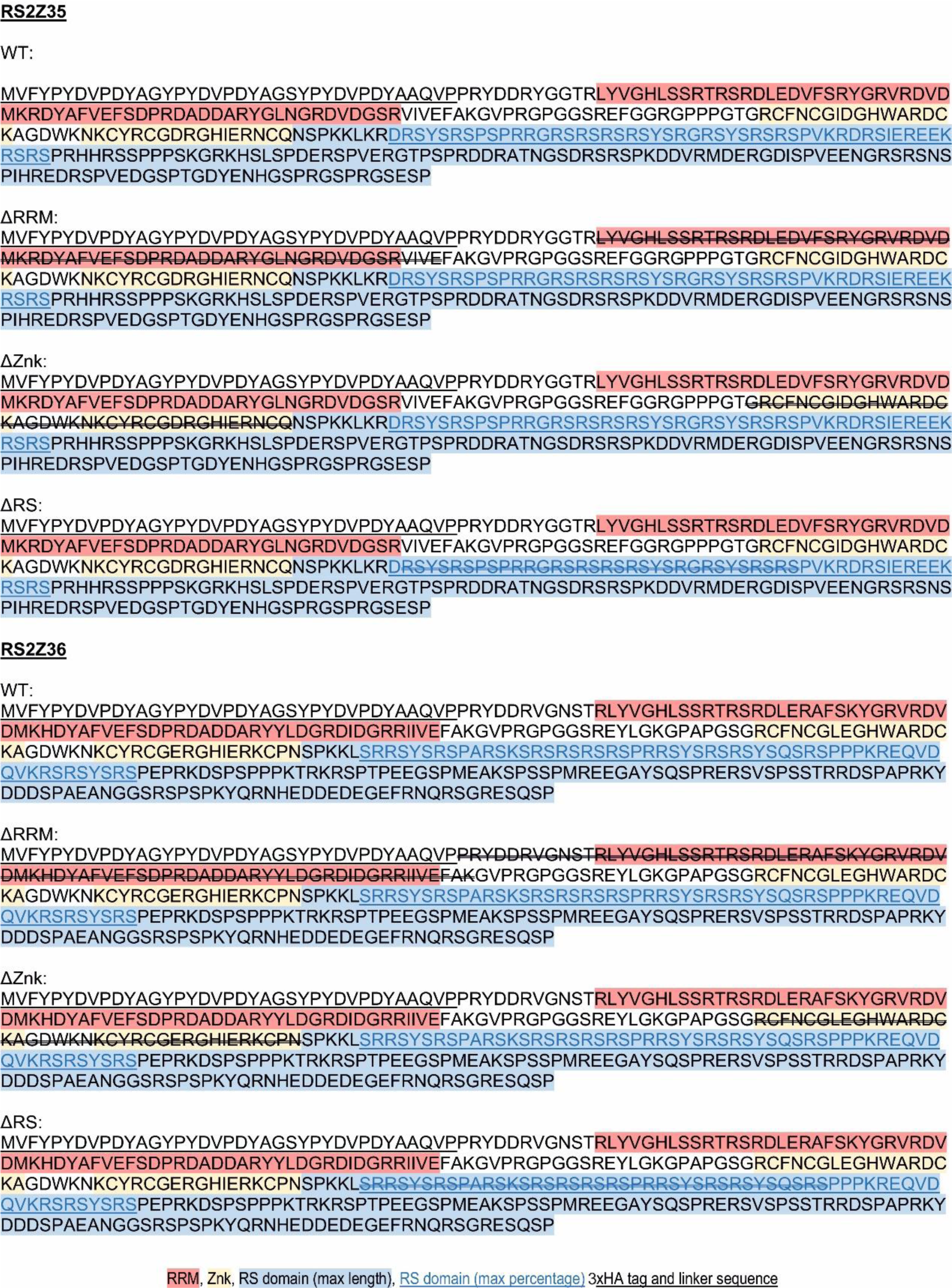
Sequences of deletion mutants. This figure is Supplementary to Fig. 2d. Red color indicates the RRM, yellow the ZnKs and blue the C-terminal with the RS-rich region shown underlined. The underlined N-terminal region indicated the 3xHA tag.

**Supplementary Figure S4.**
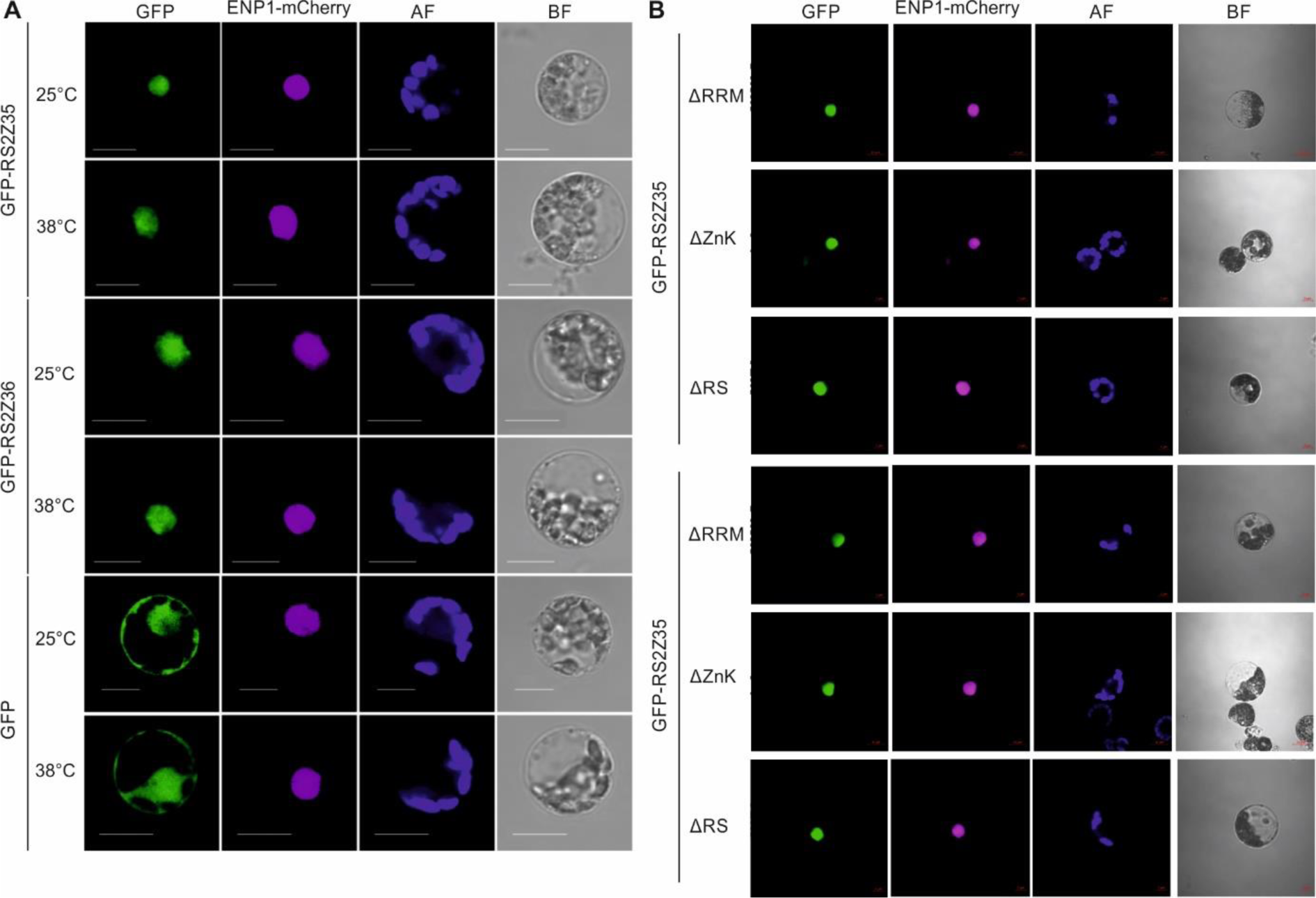
Localization of GFP-tagged RS2Z35 and RS2Z36 domain deletion mutants in tomato protoplasts. Plasmids coding for the respective wild type (A) or mutant (B) RS2Z genes were co-transformed with plasmid expressing the nuclear marker ENP1-mCherry. A plasmid carrying an expression cassette of GFP alone was used as control (A). Protoplasts expressing GFP or the wild type proteins were either kept at 25°C or exposed for 1 hour to 38°C. Images were taken by a confocal laser scanning microscope. AF: autofluorescence; BF: Bright field.

**Supplementary Figure S5.**
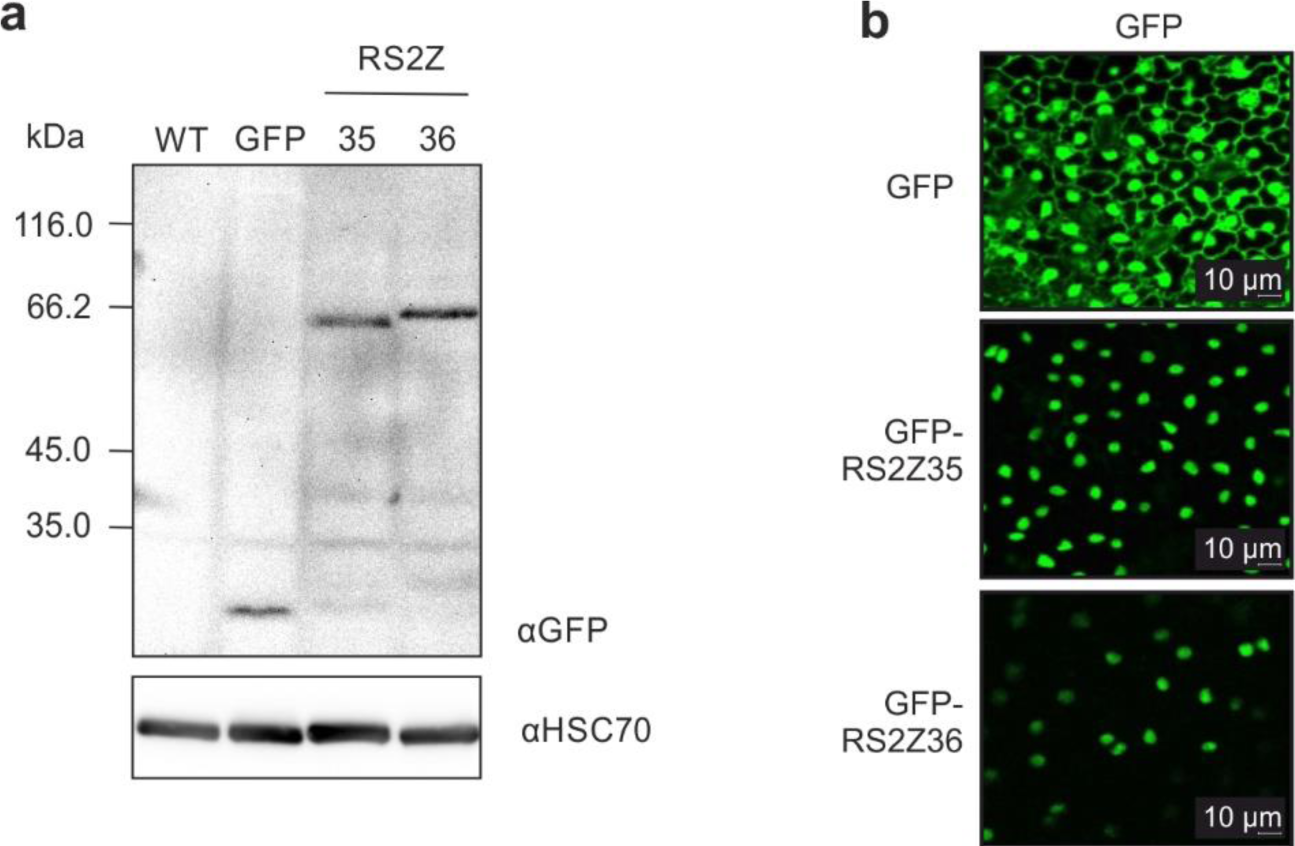
Expression of GFP and GFP-RS2Z35, GFP-RS2Z36 constructs in leaves of transgenic tomato plants. (a) Immunoblot analysis of GFP and GFP-RS2Z proteins in the respective transgenic lines or WT plants. HSC70 is shown as loading control. (b) Localization of GFP or GFP-RS2Z proteins in cells of the adaxial site of leaves in the corresponding transgenic lines based on confocal laser scanning microscopy based on the GFP signal.

**Supplementary Figure S6.**
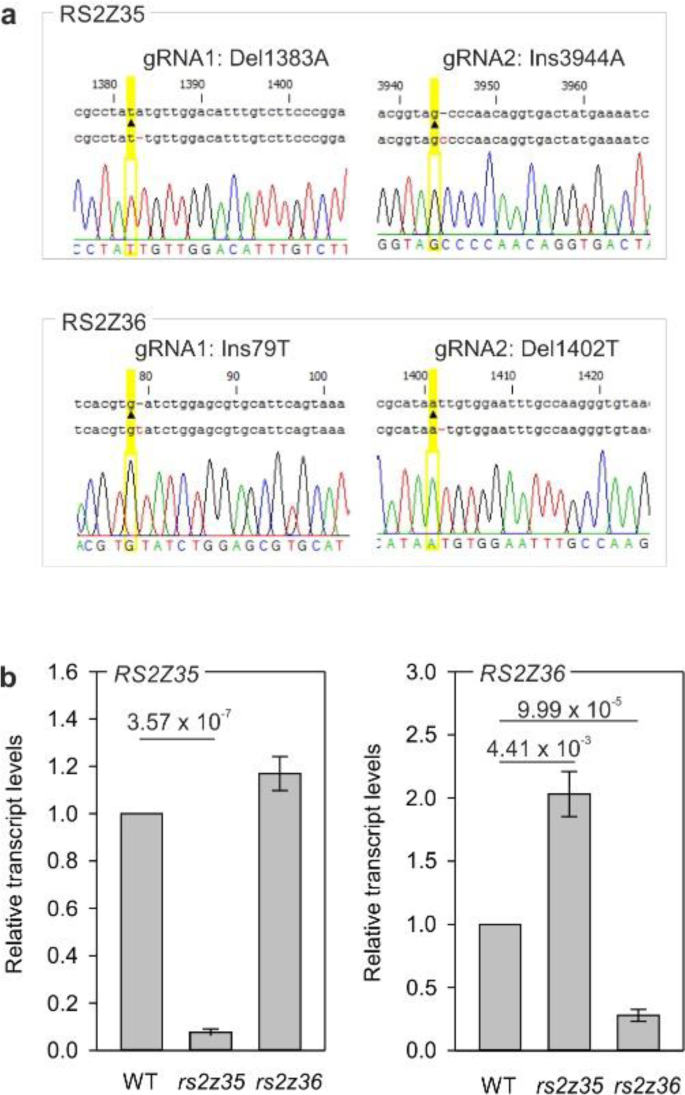
CRISPR-Cas9 mutation of RS2Z genes. (a) Sequence of mutations in RS2Z35 and RS2Z36 genes. The position of the mutation is indicated in relation to the start nucleotide of the gene (based on ITAG4.0 annotation). The electropherograms showing the Sanger sequencing analysis was done in T3 generation in T-DNA free plants, carrying RS2Z homozygous mutations. (b) Relative transcript levels of *RS2Z35* and *RS2Z36* genes in the two RS2Z mutants compared to WT based on qRT-PCR. Error bars indicate standard deviation from 3 independent biological replicated. P-values are indicated on top, and correspond to pairwise t-test analysis. EF1a was used a reference gene for normalization. This is Supplementary to Figure 3.

**Supplementary Figure S7.**
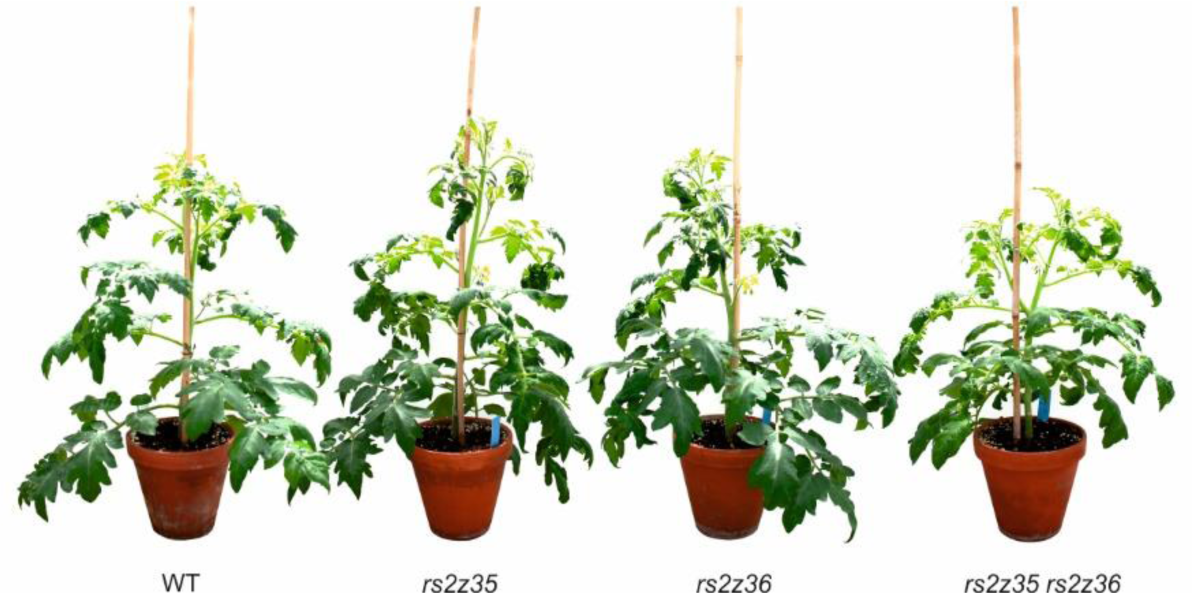
Approximately 12-week old wild type, single and double *rs2z* mutants grown in the greenhouse (25°C / 16 h light (120 μS light intensity on the canopy) / 22°C / 8 h dark).

**Supplementary Figure S8.**
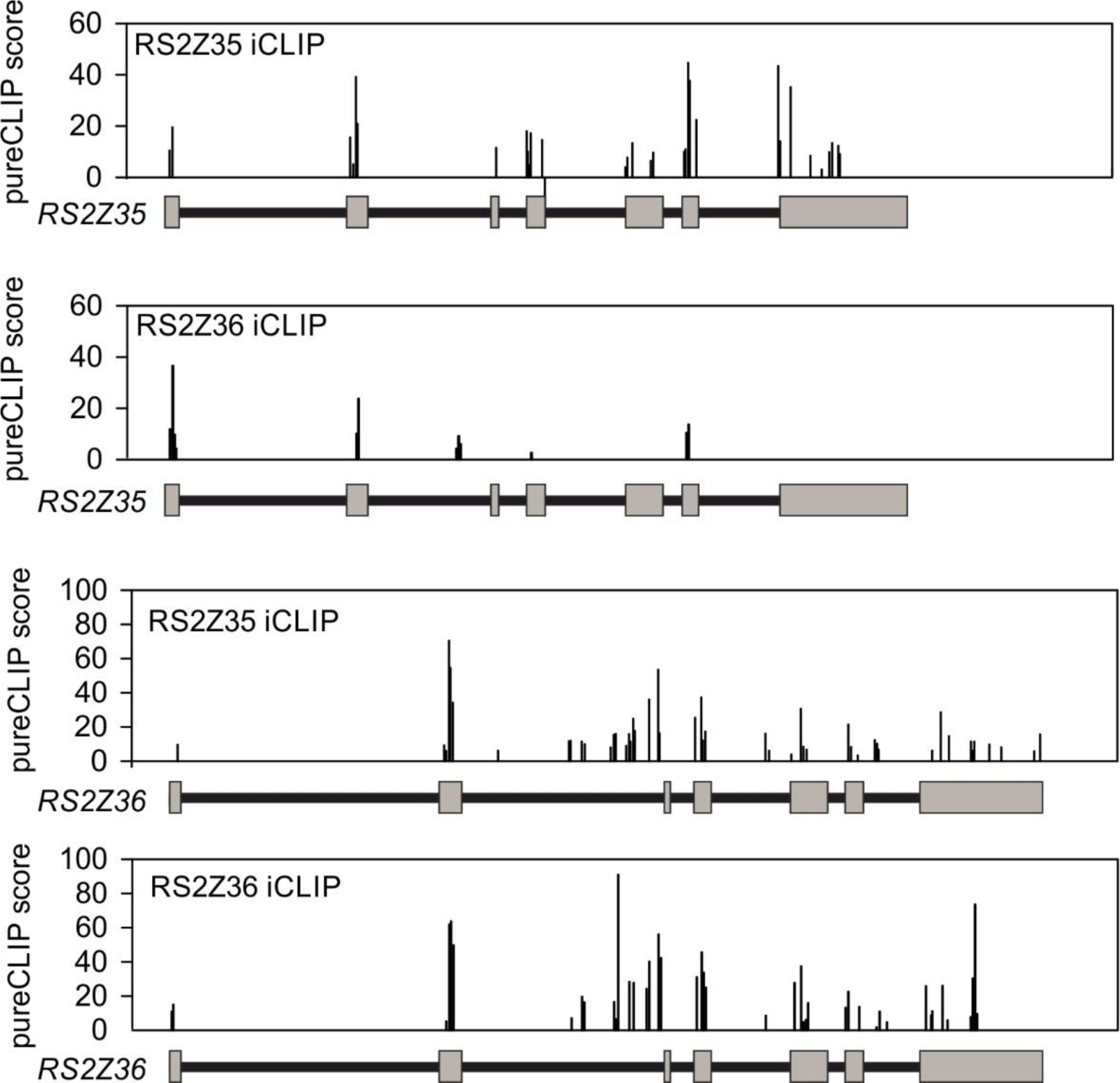
Binding sites of RS2Z35 and RS2Z36 on the two genes. For each gene the exon/intron structure is indicated below. Y-scales show the PureCLIP score for each binding site.

**Supplementary Figure S9.**
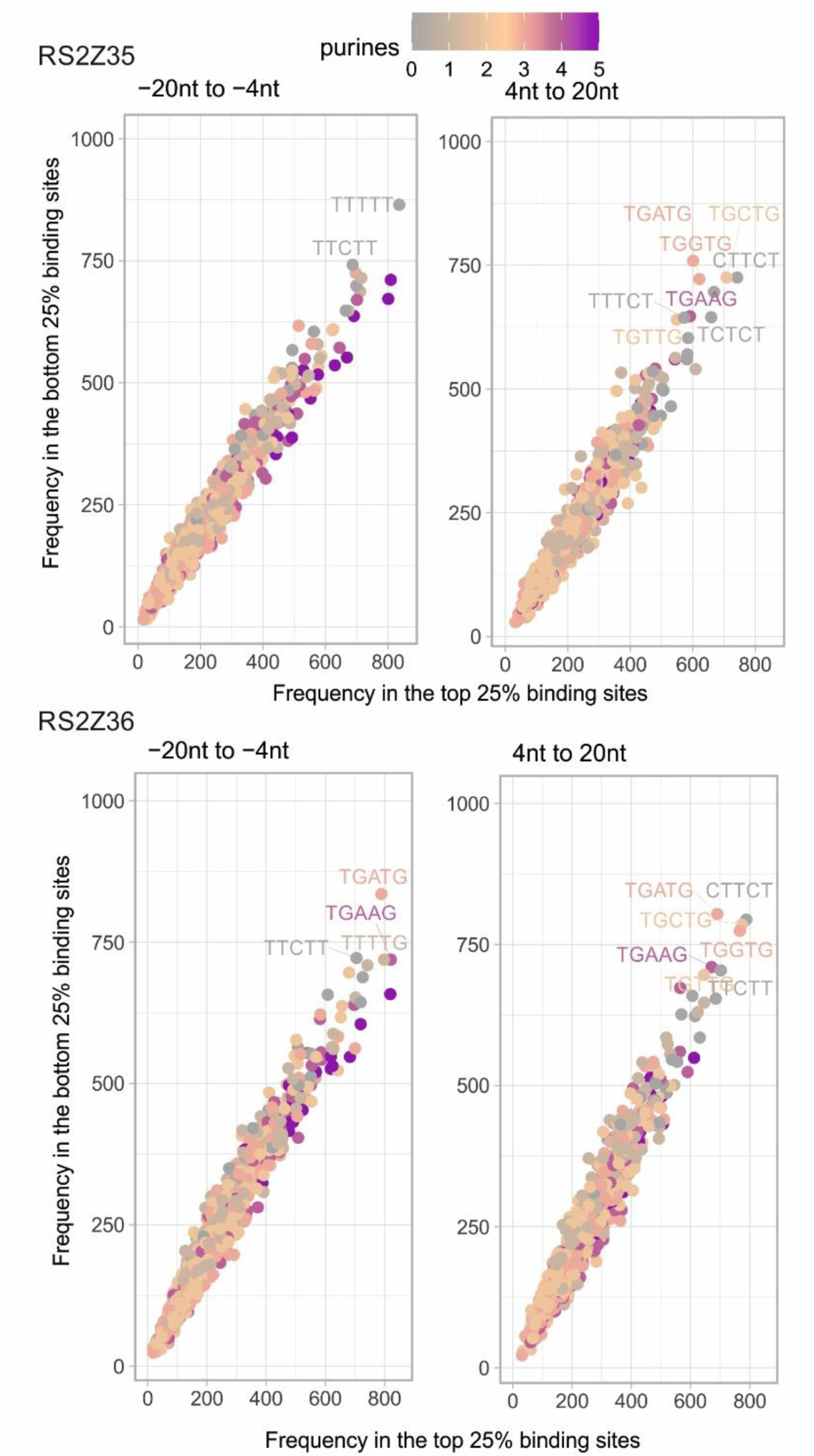
Plots of all pentamers in a 17-nt window upstream (left, −20 nt to −4 nt) or downstream (right, 4 nt to 20 nt) of the top 25% and bottom 25% binding sites (based on the PureCLIP score as a proxy for binding site strength). The color of indicates the number of purines in the pentamers. This is Supplementary to Fig. 5i.

**Supplementary Figure S10.**
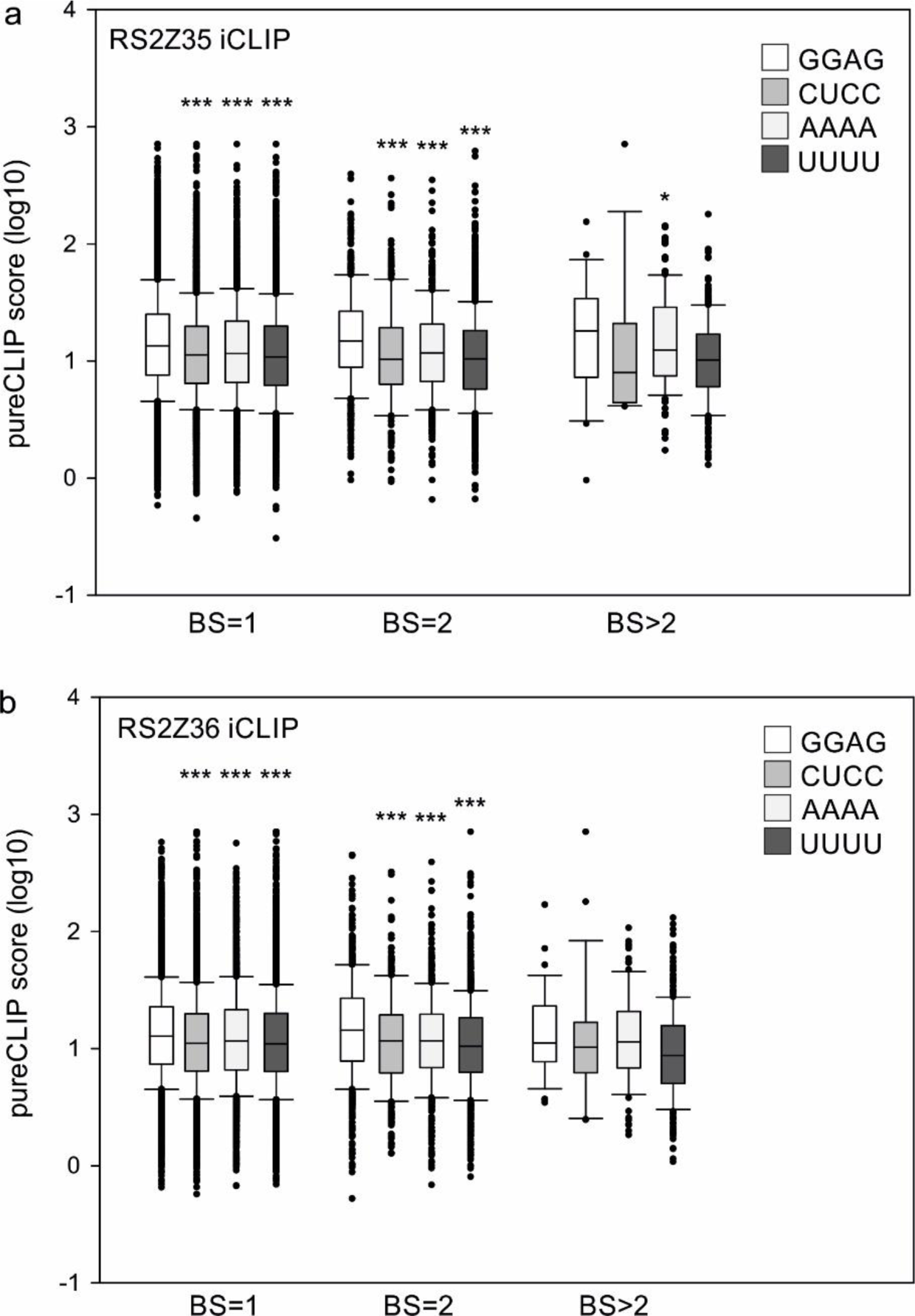
Binding strength of RS2Z35 and RS2Z36 in presence of one, two or at least three GGAG motifs within a 15 nucleotide window. Binding strength is shown as pureCLIP score (log10). CUCC, AAAA and UUUU are shown as controls. Raw values can be found in Supplementary Dataset 9. Asterisks indicate statistically significant difference between one of control motifs and GGAG, based on ANOVA and Duncan’s Multiple Range tests (* *p* < 0.05; *** *p* < 0.001).

